# NmeCas9 is an intrinsically high-fidelity genome editing platform

**DOI:** 10.1101/172650

**Authors:** Nadia Amrani, Xin D. Gao, Pengpeng Liu, Alireza Edraki, Aamir Mir, Raed Ibraheim, Ankit Gupta, Kanae E. Sasaki, Tong Wu, Paul D. Donohoue, Alexander H. Settle, Alexandra M. Lied, Kyle McGovern, Chris K. Fuller, Peter Cameron, Thomas G. Fazzio, Lihua Julie Zhu, Scot A. Wolfe, Erik J. Sontheimer

**Affiliations:** RNA Therapeutics Institute, University of Massachusetts Medical School, 368 Plantation Street, Worcester, MA 01605, U.S.A.; Program in Molecular Medicine, University of Massachusetts Medical School, 368 Plantation Street, Worcester, MA 01605, U.S.A.; Department of Molecular, Cell and Cancer Biology, University of Massachusetts Medical School, 368 Plantation Street, Worcester, MA 01605, U.S.A.; Department of Biochemistry and Molecular Pharmacology, University of Massachusetts Medical School, 368 Plantation Street, Worcester, MA 01605, U.S.A.; Program in Bioinformatics and Integrative Biology, University of Massachusetts Medical School, 368 Plantation Street, Worcester, MA 01605, U.S.A.; Caribou Biosciences, Inc., 2929 7^th^ Street, Suite 105, Berkeley, CA 94710, U.S.A.

**Keywords:** Cas9, CRISPR, sgRNA, Protospacer adjacent motif, off-target, *Neisseria meningitidis*

## Abstract

**Background:** The development of CRISPR genome editing has transformed biomedical research. Most applications reported thus far rely upon the Cas9 protein from *Streptococcus pyogenes* SF370 (SpyCas9). With many RNA guides, wild-type SpyCas9 can induce significant levels of unintended mutations at near-cognate sites, necessitating substantial efforts toward the development of strategies to minimize off-target activity. Although the genome-editing potential of thousands of other Cas9 orthologs remains largely untapped, it is not known how many will require similarly extensive engineering to achieve single-site accuracy within large (e.g. mammalian) genomes. In addition to its off-targeting propensity, SpyCas9 is encoded by a relatively large (~4.2 kb) open reading frame, limiting its utility in applications that require size-restricted delivery strategies such as adeno-associated virus vectors. In contrast, some genome-editing-validated Cas9 orthologs (e.g. from *Staphylococcus aureus, Campylobacter jejuni, Geobacillus stearothermophilus* and *Neisseria meningitidis*) are considerably smaller and therefore better suited for viral delivery.

**Results:** Here we show that wild-type NmeCas9, when programmed with guide sequences of natural length (24 nucleotides), exhibits a nearly complete absence of unintended editing in human cells, even when targeting sites that are prone to off-target activity with wildtype SpyCas9. We also validate at least six variant protospacer adjacent motifs (PAMs), in addition to the preferred consensus PAM (5’-N_4_GATT-3’), for NmeCas9 genome editing in human cells.

**Conclusions:** Our results show that NmeCas9 is a naturally high-fidelity genome editing enzyme and suggest that additional Cas9 orthologs may prove to exhibit similarly high accuracy, even without extensive engineering.

## BACKGROUND

Over the past decade, clustered, regularly interspaced, short palindromic repeats (CRISPRs) have been revealed as genomic sources of small RNAs (CRISPR RNAs, crRNAs) that specify genetic interference in many bacteria and most archaea [1–3]. CRISPR sequences include “spacers,” which often match sequences of previously encountered invasive nucleic acids such as phage genomes and plasmids.

In conjunction with CRISPR-associated (Cas) proteins, crRNAs recognize target nucleic acids (DNA, RNA, or both, depending on the system) by base pairing, leading to their destruction. The primary natural function of CRISPR-Cas systems is to provide adaptive immunity against phages [4, 5] and other mobile genetic elements [6]. CRISPR-Cas systems are divided into two main classes: Class 1, with large, multi-subunit effector complexes, and Class 2, with single-protein-subunit effectors [7]. Both CRISPR-Cas classes include multiple types based primarily on the identity of a signature effector protein. Within Class 2, the Type II systems are the most abundant and the best characterized. The interference function of Type II CRISPR-Cas systems requires the Cas9 protein, the crRNA, and a separate non-coding RNA known as the trans-activating crRNA (tracrRNA) [8–10]. Successful interference also requires that the DNA target (the “protospacer”) be highly complementary to the spacer portion of the crRNA, and that the PAM consensus be present at neighboring base pairs [11, 12].

Following the discovery that Type II interference occurs via double-strand breaks (DSBs) in the DNA target [9], the Cas9 protein was shown to be the only Cas protein required for Type II interference, to be manually reprogrammable via engineered CRISPR spacers, and to be functionally portable between species that diverged billions of years ago [10]. Biochemical analyses with purified Cas9 confirmed its role as a crRNA-guided, programmable nuclease that induces R-loop formation between the crRNA and one dsDNA strand, and that cleaves the crRNA-complementary and noncomplementary strands with its HNH and RuvC domains, respectively [13, 14]. *In vitro* cleavage reactions also showed that the tracrRNA is essential for DNA cleavage activity, and that the naturally separate crRNA and tracrRNA could retain function when fused into a single-guide RNA (sgRNA) [14]. Several independent reports then showed that the established DSB-inducing activity of Cas9 could be elicited not only *in vitro* but also in living cells, both bacterial [15] and eukaryotic [16–20]. As with earlier DSB-inducing systems [21], cellular repair of Cas9-generated DSBs by either non-homologous end joining (NHEJ) or homology-directed repair (HDR) enabled live-cell targeted mutagenesis, and the CRISPR-Cas9 system has now been widely adopted as a facile genome-editing platform in a wide range of organisms [22–24]. In addition to genome editing, catalytically inactivated Cas9 (“dead” Cas9, dCas9) retains its sgRNA-guided DNA binding function, enabling fused or tethered functionalities to be delivered to precise genomic loci [25, 26]. Similar RNA-guided tools for genome manipulations have since been developed from Type V CRISPR-Cas systems that use the Cas12a (formerly Cpf1) enzyme [27].

Type II CRISPR-Cas systems are currently grouped into three subtypes (II-A, II-B and II-C) [7, 28]. The vast majority of Cas9 characterization has been done on a single Type II-A ortholog, SpyCas9, in part due to its consistently high genome editing activity. SpyCas9’s sgRNAs typically contain a 20-nt guide sequence (the spacer-derived sequence that base pairs to the DNA target [8, 14]). The PAM requirement for SpyCas9 is 5’-NGG-3’ (or, less favorably, 5’-NAG-3’), after the 3’ end of the protospacer’s crRNA-noncomplementary strand [8, 14]. Based on these and other parameters, many sgRNAs directed against potentially targetable sites in a large eukaryotic genome also have near-cognate sites available to it that lead to unintended, “off-target” editing. Indeed, off-target activity by SpyCas9 has been well-documented with many sgRNA-target combinations [29, 30], prompting the development of numerous approaches to limit editing activity at unwanted sites [31–36]. Although these strategies have been shown to minimize off-targeting to various degrees, they do not always abolish it, and they can also reduce on-target activity, at least with some sgRNAs. Furthermore, each of these approaches has required extensive testing, validation, and optimization, and in some cases [33, 37, 38] depended heavily upon prior high-resolution structural characterization [39–42].

Thousands of other Cas9 orthologs have been documented [7, 28, 43, 44], providing tremendous untapped potential for additional genome editing capabilities beyond those offered by SpyCas9. Many Cas9 orthologs will provide distinct PAM specificities, increasing the number of targetable sites in any given genome. Many pair-wise Cas9 combinations also have orthogonal guides that load into one ortholog but not the other, facilitating multiplexed applications [44–46]. Finally, some Cas9 orthologs (especially those from subtype II-C) are hundreds of amino acids smaller than the 1,368 amino acid SpyCas9 [7, 43, 44], and are therefore more amenable to combined Cas9/sgRNA delivery via a single size-restricted vector such as adeno-associated virus (AAV) [47, 48]. Finally, there may be Cas9 orthologs that exhibit additional advantages such as greater efficiency, natural hyper-accuracy, distinct activities, reduced immunogenicity, or novel means of control over editing. Deeper exploration of the Cas9 population could therefore enable expanded or improved genome engineering capabilities.

We have used *N. meningitidis* (strain 8013) as a model system for the interference functions and mechanisms of Type II-C CRISPR-Cas systems [49–52]. In addition, we and others previously reported that the Type II-C Cas9 ortholog from *N. meningitidis* (NmeCas9) can be applied as a genome engineering platform [46, 53, 54]. At 1,082 amino acids, NmeCas9 is 286 residues smaller than SpyCas9, making it nearly as compact as SauCas9 (1,053 amino acids) and well within range of all-in-one AAV delivery. Its spacer-derived guide sequences are longer (24 nts) than those of most other Cas9 orthologs [51], and like SpyCas9, it cleaves both DNA strands between the third and fourth nucleotides of the protospacer (counting from the PAM-proximal end). NmeCas9 also has a longer PAM consensus (5’-N_4_GATT-3’, after the 3’ end of the protospacer’s crRNA-noncomplementary strand) [44, 46, 51–54], leading to a lower density of targetable sites compared to SpyCas9. Considerable variation from this consensus is permitted during bacterial interference [46, 52], and a smaller number of variant PAMs can also support targeting in mammalian cells [53, 54]. Unlike SpyCas9, NmeCas9 has been found to cleave the DNA strand of RNA-DNA hybrid duplexes in a PAM-independent fashion [52, 55], and can also catalyze PAM-independent, spacer-directed cleavage of RNA [56]. Recently, natural Cas9 inhibitors (encoded by bacterial mobile elements) have been identified and validated in *N. meningitidis* and other bacteria with type II-C systems, providing for genetically encodable off-switches for NmeCas9 genome editing [57, 58]. These “anti-CRISPR” (Acr) proteins [59] enable temporal, spatial, or conditional control over the NmeCas9 system. Natural inhibitors of Type II-A systems have also been discovered in *Listeria monocytogenes* [60] and *Streptococcus thermophilus* [61], some of which are effective at inhibiting SpyCas9.

The longer PAM consensus, longer guide sequence, or enzymological properties of NmeCas9 could result in a reduced propensity for off-targeting, and targeted deep sequencing at bioinformatically predicted near-cognate sites is consistent with this possibility [54]. A high degree of genome-wide specificity has also been noted for the dNmeCas9 platform [62]. However, the true, unbiased accuracy of NmeCas9 is not known, since empirical assessments of genome-wide off-target editing activity (independent of bioinformatics prediction) have not been reported for this ortholog. Here we define and confirm many of the parameters of NmeCas9 editing activity in mammalian cells including PAM sequence preferences, guide length limitations, and off-target profiles. Most notably, we use two empirical approaches (GUIDE-seq [63] and SITE-Seq [64] to define NmeCas9 off-target profiles and find that wild-type NmeCas9 is a high-fidelity genome editing platform in mammalian cells, with far lower levels of off-targeting than wild-type SpyCas9. These results further validate NmeCas9 as a genome engineering platform, and suggest that continued exploration of Cas9 orthologs could identify additional RNA-guided nucleases that exhibit favorable properties, even without the extensive engineering efforts that have been applied to SpyCas9 [31, 34, 35].

## RESULTS

### Co-expressed sgRNA increases NmeCas9 accumulation in mammalian cells

Previously we demonstrated that NmeCas9 (derived from *N. meningitidis* strain 8013 [51]) can efficiently edit chromosomal loci in human stem cells using either dual RNAs (crRNA + tracrRNA) or a sgRNA [53]. To further define the efficacy and requirements of NmeCas9 in mammalian cells, we first constructed an all-in-one plasmid (pEJS15) that delivers both NmeCas9 protein and a sgRNA in a single transfection vector, similar to our previous all-in-one dual-RNA plasmid (pSimple-Cas9-Tracr-crRNA; Addgene #47868) [53]. The pEJS15 plasmid expresses NmeCas9 fused to a C-terminal single-HA epitope tag and nuclear localization signal (NLS) sequences at both N- and C-termini under the control of the elongation factor-1α (EF1α) promoter. The sgRNA cassette (driven by the U6 promoter) includes two *BsmBI* restriction sites that are used to clone a spacer of interest from short, synthetic oligonucleotide duplexes. First, we cloned three different bacterial spacers (spacers 9, 24 and 25) from the endogenous *N. meningitidis* CRISPR locus (strain 8013) [51, 52] to express sgRNAs that target protospacer (ps) 9, ps24 or ps25, respectively (Supplemental Fig. 1A). None of these protospacers have cognate targets in the human genome. We also cloned a spacer sequence to target an endogenous genomic NmeCas9 target site (NTS) from chromosome 10 that we called N-TS3 (Table 1). Two of the resulting all-in-one plasmids (spacer9/sgRNA and N-TS3/sgRNA), as well as a plasmid lacking the sgRNA cassette, were transiently transfected into HEK293T cells for 48 hours, and NmeCas9 expression was assessed by anti-HA western blot (Fig. 1A). As a positive control we also included a sample transfected with a SpyCas9-expressing plasmid (triple-HA epitope-tagged, and driven by the cytomegalovirus (CMV) promoter) [65] (Addgene #69220). Full-length NmeCas9 was efficiently expressed in the presence of both sgRNAs (lanes 3 and 4). However, the abundance of the protein was much lower in the absence of sgRNA (lane 2). A different Type II-C Cas9 (CdiCas9) was shown to be dramatically stabilized by its cognate sgRNA when subjected to proteolysis *in vitro* [55]; if similar resistance to proteolysis occurs with NmeCas9 upon sgRNA binding, it could explain some or all of the sgRNA-dependent increase in cellular accumulation.

**Figure 1.**
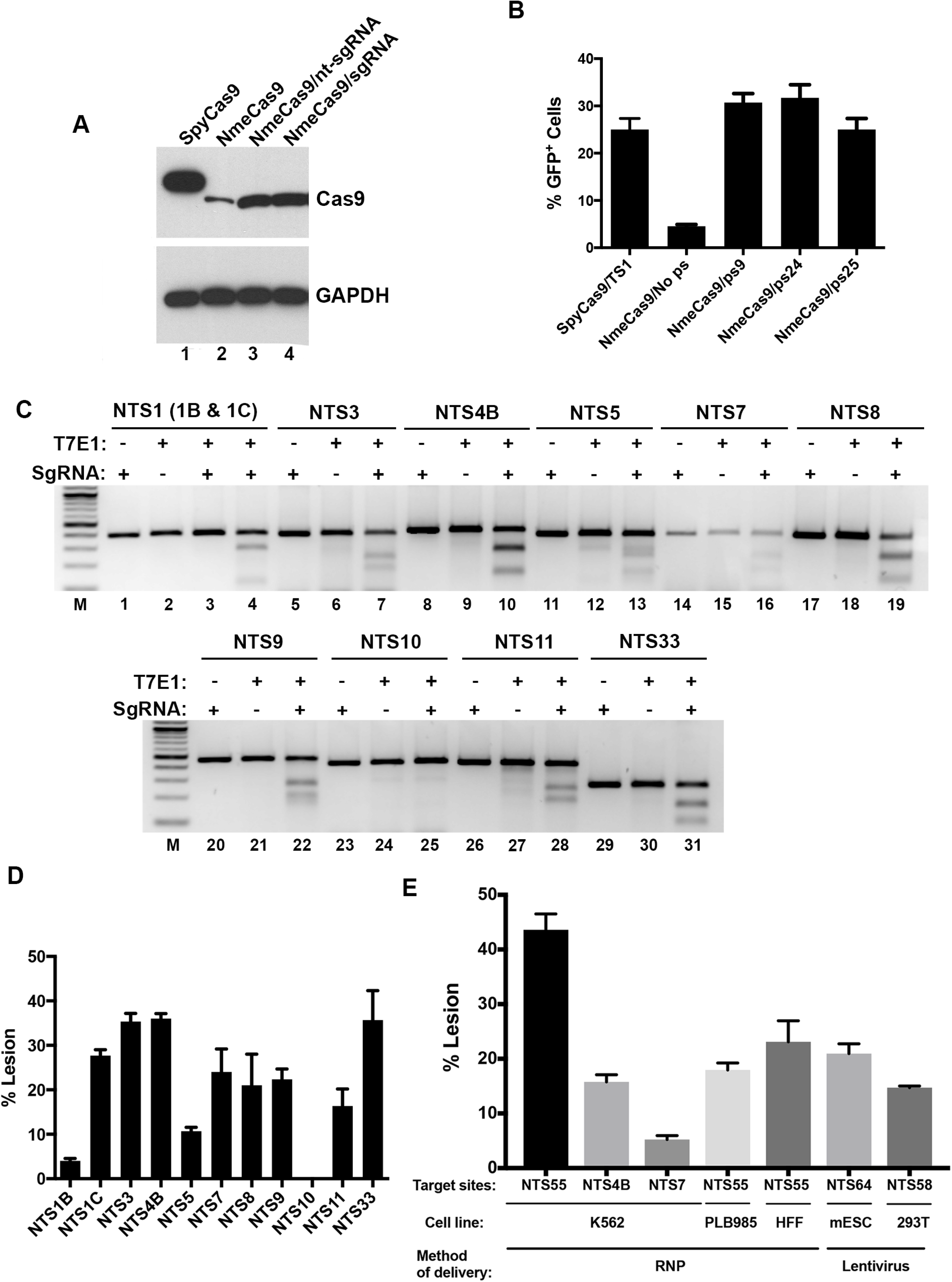
NmeCas9 expression and activity in human (HEK293T) cells. (*A*) Western blot detection of HA-tagged NmeCas9 in transiently transfected HEK293T cells. Lane 1: Cells transfected with SpyCas9 plasmid under the control of the CMV promoter. Lane 2: Cells transfected with NmeCas9 plasmid under the control of the elongation factor-1α (EF1α) promoter. Lane 3: Cells expressing NmeCas9 and a non-targeting sgRNA (nt-sgRNA), which lacks a complementary site in the human genome. Lane 4: Cells expressing NmeCas9 and a sgRNA targeting chromosomal site NTS3. Upper panel: Anti-HA western blot. Lower panel: Anti-GAPDH western blot as a loading control. (*B*) NmeCas9 targeting co-transfected split-GFP reporter with ps9, ps24 and ps25 sites. Plasmid cleavage by SpyCas9 is used as a positive control, and a reporter without a guide-complementary site (No ps: no protospacer) is used as a negative control to define background levels of recombination leading to GFP+ cells. (*C*) NmeCas9 programmed independently with different sgRNAs targeting eleven genomic sites flanked by an N4GATT PAM, detected by T7E1 analysis. (*D*) Quantitation of editing efficiencies from three independent biological replicates performed on different days. Error bars indicate ± standard error of the mean (± s.e.m.). (*E*) Genomic lesions with NmeCas9 programmed independently with different guides in different cell lines and using different methods of delivery.

**Table 1.**
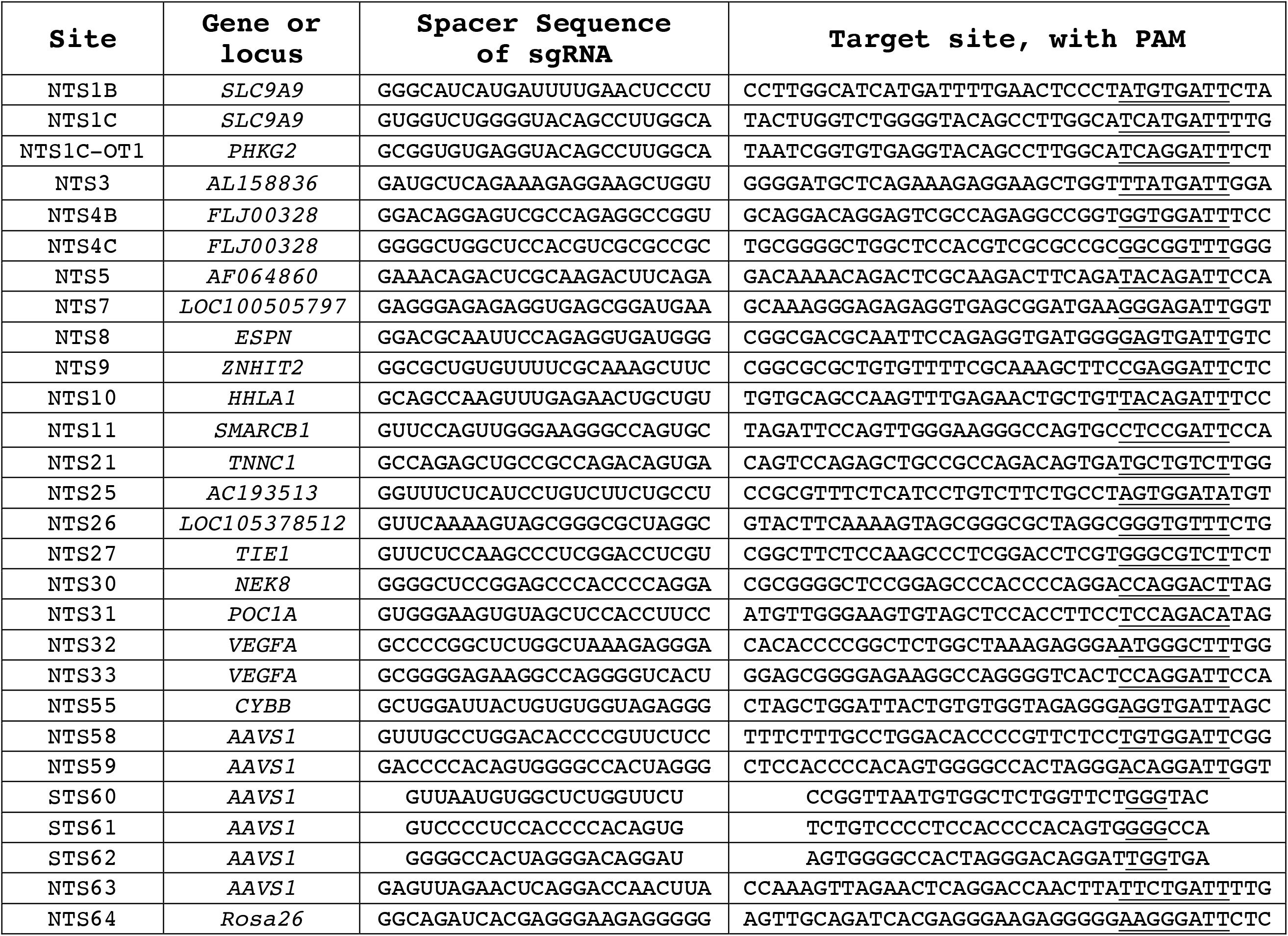
NmeCas9 or SpyCas9 guide and target sequences used in this study. NTS, NmeCas9 target site; STS, SpyCas9 target site. The sgRNA spacer sequences (5’➔3’) are shown with their canonical lengths, and with a 5’-terminal G residue; non-canonical lengths are described in the text and figures. Target site sequences are also 5’➔3’ and correspond to the DNA strand that is non-complementary to the sgRNA, with PAM sequences underlined.

### Efficient editing in mammalian cells by NmeCas9

To establish an efficient test system for NmeCas9 activity in mammalian cells, we used a co-transfected fluorescent reporter carrying two truncated, partially overlapping GFP fragments that are separated by a cloning site [66] into which we can insert target protospacers for NmeCas9. Cleavage promotes a single-strand-annealing-based repair pathway that generates an intact GFP open reading frame (ORF), leading to fluorescence [66] that can be scored after 48 hours by flow cytometry. We generated reporters carrying three validated bacterial protospacers (ps9, ps24 and ps25, as described above) [51, 52] for transient cotransfection into HEK293T cells along with the corresponding NmeCas9/sgRNA constructs. Figure 1B shows that all three natural protospacers of NmeCas9 can be edited in human cells and the efficiency of GFP induction was comparable to that observed with SpyCas9 (Fig. 1B).

Next, we reprogrammed NmeCas9 by replacing the bacterially-derived spacers with a series of spacers designed to target eleven human chromosomal sites with an N4GATT PAM (Table 1). These sgRNAs induced insertion/deletion (indel) mutations at all sites tested, except NTS10 (Fig. 1C, lanes 23-25), as determined by T7 Endonuclease 1 (T7E1) digestion (Fig. 1C). The editing efficiencies ranged from 5% for NTS1B site to 47% in the case of NTS33 (Fig. 1D), though T7E1 tends to underestimate the true frequencies of indel formation [67]. These data confirm that NmeCas9 can induce, with variable efficiency, edits at many genomic target sites in human cells. Furthermore, we demonstrated NmeCas9 genome editing in multiple cell lines and via distinct delivery modes. Nucleofection of NmeCas9 ribonucleoprotein (RNP) (loaded with an *in vitro*-transcribed sgRNA) led to indel formation at three sites in K562 chronic myelogenous leukemia cells and in hTERT-immortalized human foreskin fibroblasts (gift from Dr. Job Dekker) (Fig. 1E). In addition, mouse embryonic stem cells (mESCs) and HEK293T cells were transduced with a lentivirus construct expressing NmeCas9. In these cells, transient transfection of plasmids expressing a sgRNA led to genome editing (Fig. 1E). Collectively, our results show that NmeCas9 can be used for genome editing in a range of human or mouse cell lines via plasmid transfection, RNP delivery, or lentiviral transduction.

### Functionality of truncated sgRNAs with NmeCas9

SpyCas9 can accommodate limited variation in the length of the guide region (normally 20 nucleotides) of its sgRNAs [68–71], and sgRNAs with modestly lengthened (22-nt) or shortened (17–18-nt) guide regions can even enhance editing specificity by reducing editing at off-target sites by a greater degree than they affect editing at the on-target site [68, 69]. To test the length dependence of the NmeCas9 guide sequence (normally 24 nucleotides; [51]) during mammalian editing, we constructed a series of sgRNAs containing 18, 19, 20, 21, 22, 23, and 24 nucleotides of complementarity to ps9 cloned into the split-GFP reporter plasmid (Supplemental Fig. 1B). All designed guides started with two guanine nucleotides (resulting in 1-2 positions of target non-complementarity at the very 5’ end of the guide) to facilitate transcription and to test the effects of extra 5’-terminal G residues, analogous to the SpyCas9 “GGN20” sgRNAs [68]. We then measured the abilities of these sgRNAs to direct NmeCas9 cleavage of the reporter in human cells. sgRNAs that have 20–23 nucleotides of target complementarity showed activities comparable to the sgRNA with the natural 24 nucleotides of complementarity, whereas sgRNAs containing 18 or 19 nucleotides of complementarity show lower activity (Fig. 2A).

**Figure 2.**
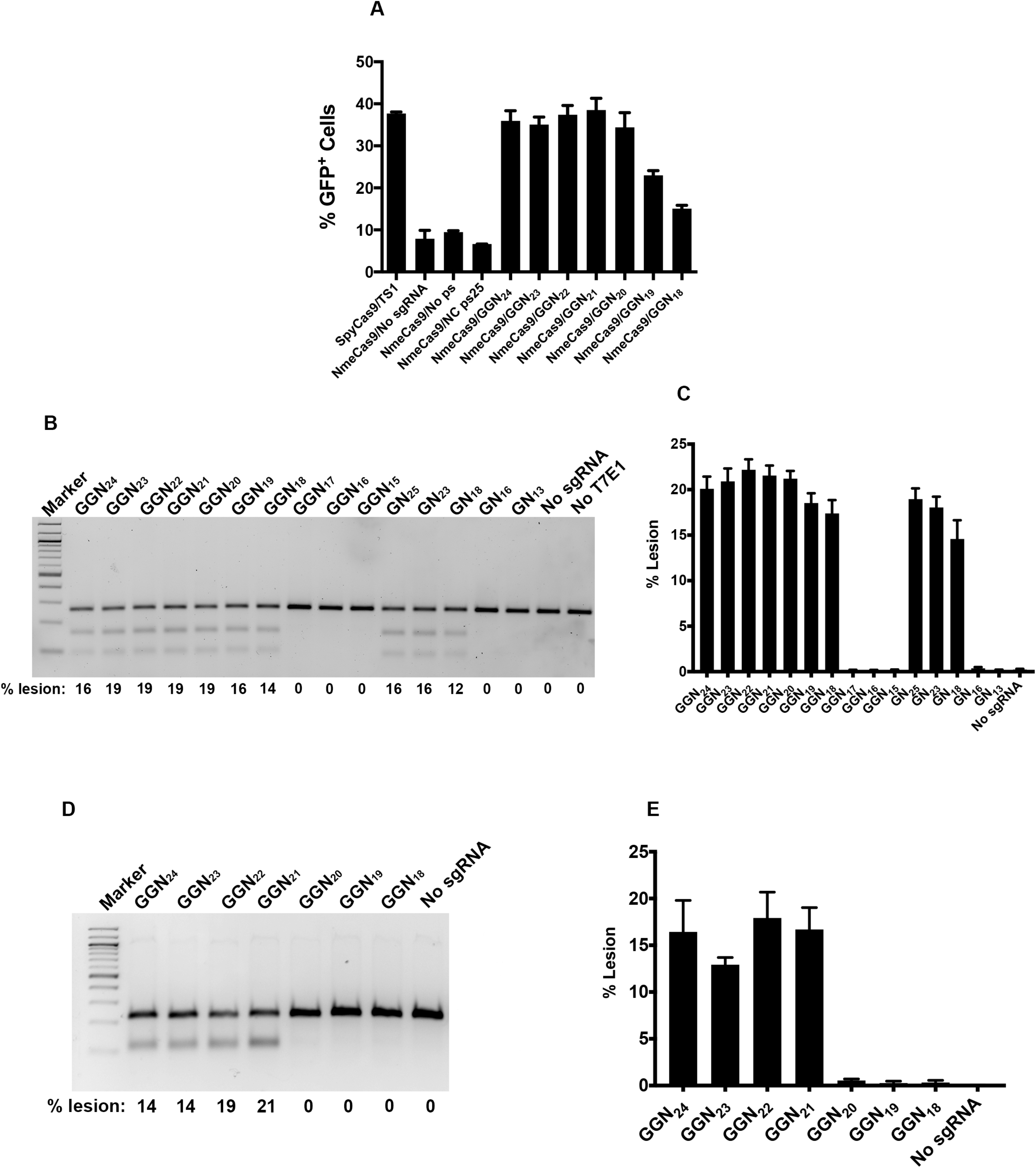
NmeCas9 guide length requirements in mammalian cells. (*A*) Split-GFP activity profile of NmeCas9 cleavage with ps9 sgRNAs bearing spacers of varying lengths (18-24 nts) along with 5’-terminal G residues to enable transcription. Bars represent mean values ± s.e.m. from three independent biological replicates performed on different days. (*B*) T7EI analysis of editing efficiencies at the NTS33 genomic target site (with an N_4_GATT PAM) with sgRNAs bearing spacers of varying lengths (13-25 nts) with 1-2 5’-terminal G residues. (*C*) Quantitation of lesion efficiencies (of experiment in *B*) from three independent biological replicates performed on different days. Error bars indicate ± standard error of the mean (± s.e.m.). (*D*) As in (*B*), but targeting the NTS32 genomic site (with an N_4_GCTT PAM). (*E*) Quantitation of lesion efficiencies (of experiment in D) from three independent biological replicates performed on different days. Error bars indicate ± standard error of the mean (± s.e.m.).

We next used a native chromosomal target site (NTS33 in *VEGFA*, as in Figs. 1C and 1D) to test the editing efficiency of NmeCas9 spacers of varying lengths (Supplemental Fig. 1C). sgRNA constructs included one or two 5’-terminal guanine residues to enable transcription by the U6 promoter, sometimes resulting in 1–2 nucleotides of target non-complementarity at the 5’ end of the guide sequence. sgRNAs with 20, 21, or 22 nucleotides of target complementarity (GGN18, GGN19, and GGN20, respectively) performed comparably to the natural guide length (24 nucleotides of complementarity, GN_23_) at this site (Fig. 2B-C), and within this range, the addition of 1–2 unpaired G residues at the 5’ end had no adverse effect. These results are consistent with the results obtained with the GFP reporter (Fig. 2A). sgRNAs with guide lengths of 19 nucleotides or shorter, along with a single mismatch in the first or second position (GGN17, GGN16, and GGN15), did not direct detectable editing, nor did a sgRNA with perfectly matched guide sequences of 17 or 14 nucleotides (GN16 and GN13, respectively) (Fig. 2B-C). However, a 19-nt guide with no mismatches (GN18) successfully directed editing, albeit with slightly reduced efficiency. These results indicate that 19–26-nt guides can be tolerated by NmeCas9, but that activity can be compromised by guide truncations from the natural length of 24 nucleotides down to 17–18 nucleotides and smaller, and that single mismatches (even at or near the 5’-terminus of the guide) can be discriminated against with a 19-nt guide.

The target sites tested in Figs. 2A and 2B-C are both associated with a canonical N_4_GATT PAM, but efficient NmeCas9 editing at mammalian chromosomal sites associated with N_4_GCTT [53] and other variant PAMs [[54]; also see below] has also been reported. To examine length dependence at a site with a variant PAM, we varied guide sequence length at the N_4_GCTT-associated NTS32 site (also in *VEGFA*). In this experiment, each of the guides had two 5’-terminal G residues, accompanied by 1–2 terminal mismatches with the target sequence (Supplemental Fig. 1D). At the NTS32 site, sgRNAs with 21–24 nucleotides of complementarity (GGN24, GGN23, GGN22, and GGN21) supported editing, but shorter guides (GGN20, GGN19, and GGN18) did not (Fig. 2D-E). We conclude that sgRNAs with 20 nucleotides of complementarity can direct editing at some sites (Fig. 2B-C) but not all (Fig. 2D-E). It is possible that this minor variation in length dependence can be affected by the presence of mismatched 5’-terminal G residues in the sgRNA, the adherence of the target to the canonical N4GATT PAM consensus, or both, but the consistency of any such relationship will require functional tests at much larger numbers of sites. Nonetheless, NmeCas9 guide truncations of 1–3 nucleotides appear to be functional in most cases, in agreement with the results of others [54].

### PAM specificity of NmeCas9 in human cells

During native CRISPR interference in bacterial cells, considerable variation in the N_4_GATT PAM consensus is tolerated: although the G1 residue (N4GATT) is strictly required, virtually all other single mutations at A2 (N_4_GATT), T3 (N_4_GATT), and T4 (N_4_GATT) retain at least partial function in licensing bacterial interference [46, 52]. In contrast, fewer NmeCas9 PAM variants have been validated during genome editing in mammalian cells [53, 54]. To gain more insight into NmeCas9 PAM flexibility and specificity in mammalian cells, and in the context of an otherwise identical target site and an invariant sgRNA, we employed the split-GFP readout of cleavage activity described above. We introduced single-nucleotide mutations at every position of the PAM sequence of ps9, as well as all double mutant combinations of the four most permissive single mutants, and then measured the ability of NmeCas9 to induce GFP fluorescence in transfected HEK293T cells. The results are shown in Fig. 3A. As expected, mutation of the G1 residue to any other base reduced editing to background levels, as defined by the control reporter that lacks a protospacer [(no ps), see Fig. 3A]. As for mutations at the A2, T3 and T4 positions, four single mutants (N4GCTT, N4GTTT, N4GACT, and N4GATA) and two double mutants (N4GTCT and N4GACA) were edited with efficiencies approaching that observed with the N4GATT PAM. Two other single mutants (N4GAGT and N4GATG), and three double mutants (N4GCCT, N_4_GCTA, and N_4_GTTA) gave intermediate or low efficiencies, and the remaining mutants tested were at or near background levels. We note that some of the minimally functional or non-functional PAMs (e.g. N_4_GAAT and N_4_GATC) in this mammalian assay fit the functional consensus sequences defined previously in *E. coli* [46].

**Figure 3.**
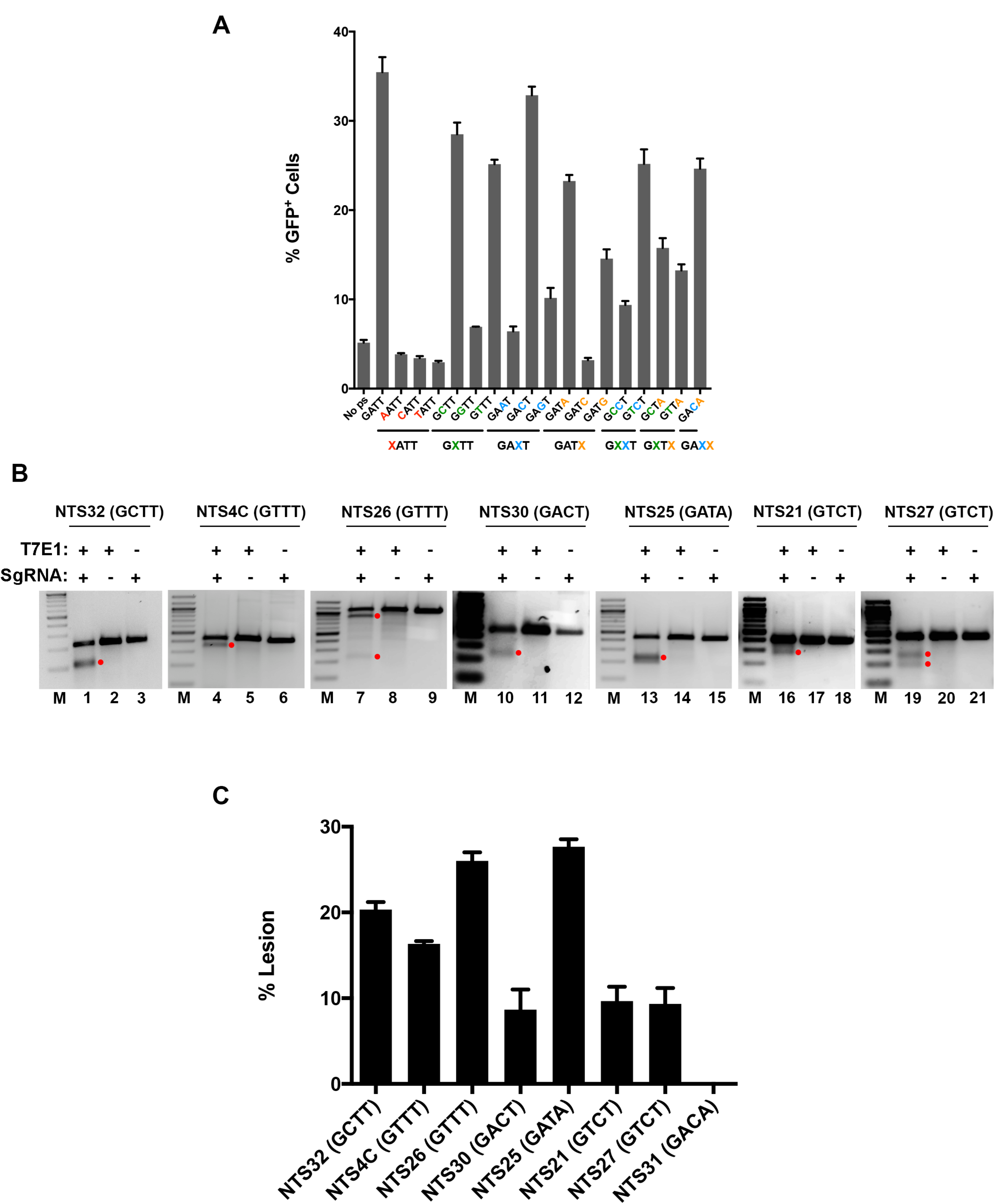
Characterization of functional PAM sequences in human (HEK293T) cells. (*A*) Split-GFP activity profile of NmeCas9 cleavage with ps9 sgRNA, with the target site flanked by different PAM sequences. Bars represent mean values ± s.e.m. from three independent biological replicates performed on different days. (*B*) T7E1 analysis of editing efficiencies at seven genomic sites flanked by PAM variants, as indicated. Products resulting from NmeCas9 genome editing are denoted by the red dots. (*C*) Quantitation of data from (*B*), as well as an additional site (NTS31; N_4_GACA PAM) that was not successfully edited. Bars represent mean values ± s.e.m. from three independent biological replicates performed on different days.

We then used T7E1 analysis to validate genome editing at eight native chromosomal sites associated with the most active PAM variants (N_4_GCTT, N_4_GTTT, N_4_GACT, N_4_GATA, N_4_GTCT, and N4GACA). Our results with this set of targets indicate that all of these PAM variants tested except N4GACA support chromosomal editing (Fig. 3B and C).

### Apo NmeCas9 is not genotoxic to mammalian cells

NmeCas9 and several other type II-C Cas9 orthologs have been shown to possess an RNA-dependent ssDNA cleavage (DNase H) activity *in vitro* [52, 55]. R-loops (regions where an RNA strand invades a DNA duplex to form a DNA:RNA hybrid, with the other DNA strand displaced) occur naturally during transcription and other cellular processes [72]. Since DNase H activity is independent of the tracrRNA or the PAM sequence, it is theoretically possible that it could degrade naturally-occurring R-loops in living cells. Global degradation of R-loops in cells could result in an increase in DNA damage detectable by increased γH2AX staining [73]. To test whether the DNase H activity of NmeCas9 could lead to an increase in γH2AX, we transduced mouse embryonic stem cells E14 (mESCs) with lentiviral plasmids expressing NmeCas9 and dNmeCas9 (which lacks DNase H activity; [52]). mESCs are ideal for this purpose as R-loops have been extensively studied in these cells and have been shown to be important for differentiation [74]. We performed γH2AX staining of these two cell lines and compared them to wildtype E14 cells. As a positive control for γH2AX induction, we exposed wildtype E14 cells to UV, a known stimulator of the global DNA damage response. Immunofluorescence microscopy of cells expressing NmeCas9 or dNmeCas9 exhibited no increase in γH2AX foci compared to wildtype E14, suggesting that sustained NmeCas9 expression is not genotoxic (Supplemental Fig. 2A). In contrast, cells exposed to UV light showed a significant increase in γH2AX levels. Flow cytometric measurements of γH2AX immunostaining confirmed these results (Supplemental Fig. 2B). These data suggest that NmeCas9 expression does not lead to a global DNA damage response in mESCs.

### Comparative analysis of NmeCas9 and SpyCas9

SpyCas9 is by far the best-characterized Cas9 orthologue, and is therefore the most informative benchmark when defining the efficiency and accuracy of other Cas9s. To facilitate comparative experiments between NmeCas9 and SpyCas9, we developed a matched Cas9 + sgRNA expression system for the two orthologs. This serves to minimize the expression differences between the two Cas9s in our comparative experiments, beyond those differences dictated by the sequence variations between the orthologues themselves. To this end, we employed the separate pCSDest2-SpyCas9-NLS-3XHA-NLS (Addgene #69220) and pLKO.1-puro-U6sgRNA-BfuA1 (Addgene #52628) plasmids reported previously for the expression of SpyCas9 (driven by the CMV promoter) and its sgRNA (driven by the U6 promoter), respectively [58, 65]. We then replaced the bacterially-derived SpyCas9 sequence (i.e., not including the terminal fusions) with that of NmeCas9 in the CMV-driven expression plasmid. This yielded an NmeCas9 expression vector (pEJS424) that is identical to that of the SpyCas9 expression vector in every way (backbone, promoters, UTRs, poly(A) signals, terminal fusions, etc.) except for the Cas9 sequence itself. Similarly, we replaced the SpyCas9 sgRNA cassette in pLKO.1-puro-U6sgRNA-BfuA1 with that of the NmeCas9 sgRNA [46, 53], yielding the NmeCas9 sgRNA expression plasmid pEJS333. This matched system facilitates direct comparisons of the two enzymes’ accumulation and activity during editing experiments. To assess relative expression levels of the identically-tagged Cas9 orthologs, the two plasmids were transiently transfected into HEK293T cells for 48 hours, and the expression of the two proteins was monitored by anti-HA western blot (Fig. 4A). Consistent with our previous data (Fig. 1A), analyses of samples from identically transfected cells show that NmeCas9 accumulation is stronger when co-expressed with its cognate sgRNA (Fig. 4A, compare lane 6 to 4 and 5), whereas SpyCas9 is not affected by the presence of its sgRNA (lanes 1-3).

**Figure 4.**
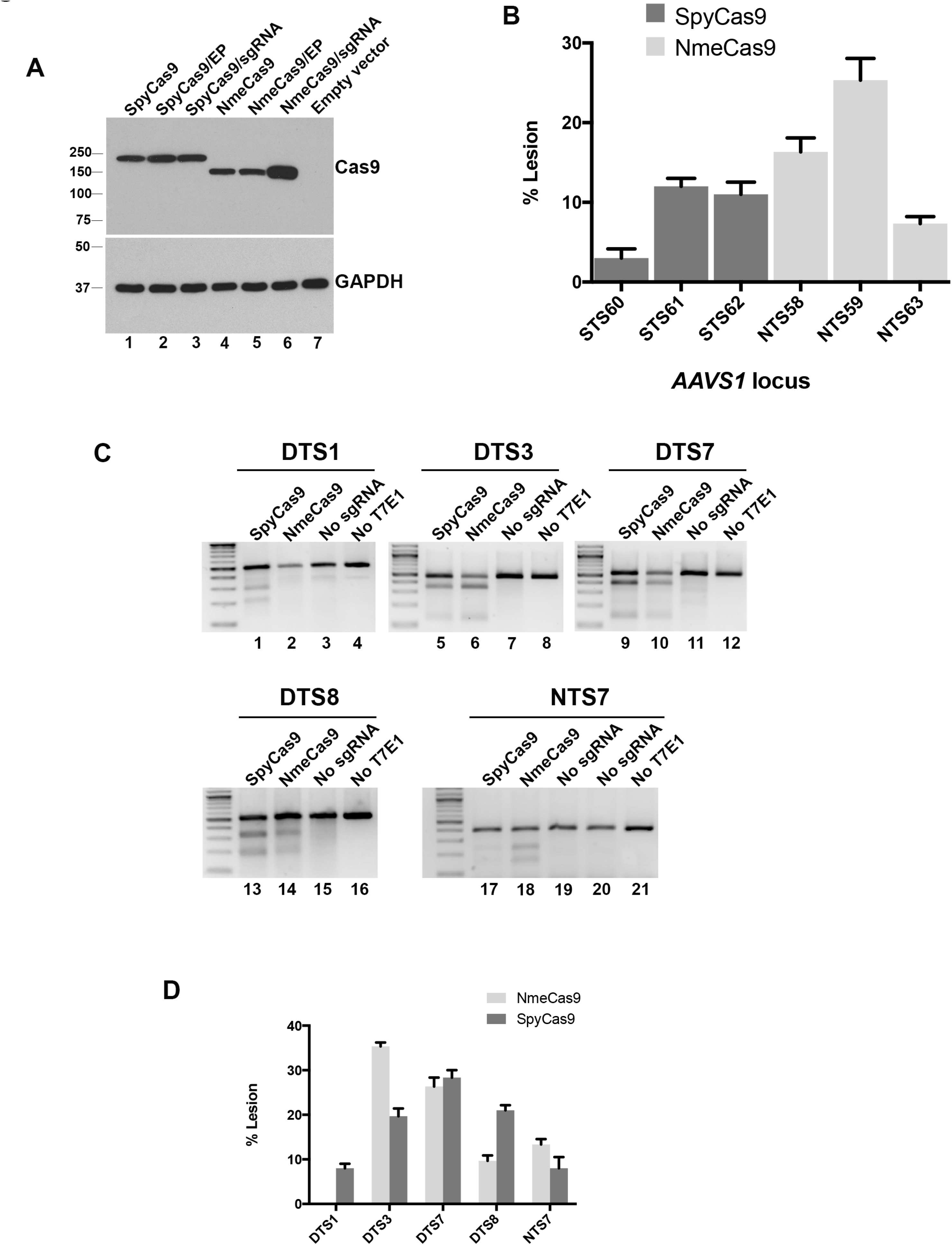
NmeCas9 and SpyCas9 have comparable editing efficiencies in human (HEK293T) cells when targeting the same chromosomal sites. (*A*) Western blot analysis of NmeCas9 and SpyCas9. HEK293T cells were transfected with the indicated Cas9 ortholog cloned in the same plasmid backbone, and fused to the same HA epitope tags and NLSs. Top panel: anti-HA western blot (EP, empty sgRNA plasmid). Bottom panel: anti-GAPDH western blot, used as a loading control. Mobilities of protein markers are indicated. (*B*) T7E1 analysis of three previously validated SpyCas9 guides targeting the *AAVS1* locus, in comparison with NmeCas9 guides targeting nearby *AAVS1* sites (mean ± s.e.m., *n* = 3). (*C*) Representative T7EI analyses comparing editing efficiencies at the dual target sites DTS1, DTS3, DTS7, DTS8, and NTS7, using the indicated Cas9/sgRNA combinations. (*D*) Qμantitation of data from (*C*)(mean ± s.e.m., *n* = 3).

For an initial comparison of the cleavage efficiencies of the two Cas9s, we chose three previously validated SpyCas9 guides targeting the *AAVS1* “safe harbor” locus [20, 75] and used the CRISPRseek package [76] to design three NmeCas9 guides targeting the same locus within a region of ~700 base pairs (Supplemental Fig. 3A). The matched Cas9/sgRNA expression systems described above were used for transient transfection of HEK293T cells. T7E1 analysis showed that the editing efficiencies were comparable, with the highest efficiency being observed when targeting the NTS59 site with NmeCas9 (Fig. 4B and Supplemental Fig. 3B).

To provide a direct comparison of editing efficiency between the SpyCas9 and NmeCas9 systems, we took advantage of the non-overlapping PAMs of SpyCas9 and NmeCas9 (NGG and N_4_GATT, respectively). Because the optimal SpyCas9 and NmeCas9 PAMs are non-overlapping, it is simple to identify chromosomal target sites that are compatible with both orthologues, i.e. that are dual target sites (DTSs) with a composite PAM sequence of NGGNGATT that is preferred by both nucleases. In this sequence context, both Cas9s will cleave the exact same internucleotide bond (N**N/N**<u>NNNGGNGATT</u>; cleaved junction in bold, and PAM region underlined), and both Cas9s will have to contend with the exact same sequence and chromatin structural context. Furthermore, if the target site contains a G residue at position −24 of the sgRNA-noncomplementary strand (relative to the PAM) and another at position −20, then the U6 promoter can be used to express perfectly-matched sgRNAs for both Cas9 orthologues. Four DTSs with these characteristics were used in this comparison (Supplemental Fig. 4A). We had previously used NmeCas9 to target a site (NTS7) that happened also to match the SpyCas9 PAM consensus, so we included it in our comparative analysis as a fifth site, even though it has a predicted rG-dT wobble pair at position −24 for the NmeCas9 sgRNA (Supplemental Fig. 4A).

We set out next to compare the editing activities of both Cas9 orthologs programmed to target the five chromosomal sites depicted in Supplemental Fig. 4A, initially via T7E1 digestion. SpyCas9 was more efficient than NmeCas9 at generating lesions at the DTS1 and DTS8 sites (Fig. 4C, lanes 1-2 and 13-14). In contrast, NmeCas9 was more efficient than SpyCas9 at the DTS3 and NTS7 sites (Fig. 4C, lanes 5-6 and 17-18). Editing at DTS7 was approximately equal with both orthologs (Fig. 4C, lanes 9-10). Data from three biological replicates of all five target sites are plotted in Fig. 4D. The remainder of our comparative studies focused on DTS3, DTS7, and DTS8, as they provided examples of target sites with NmeCas9 editing efficiencies that are greater than, equal to, or lower than those of SpyCas9, respectively. At all three of these sites, the addition of an extra 5’-terminal G residue had little to no effect on editing by either SpyCas9 or NmeCas9 (Supplemental Fig. 4B). Truncation of the three NmeCas9 guides down to 20 nucleotides (all perfectly matched) again had differential effects on editing efficiency from one site to the next, with no reduction in DTS7 editing, partial reduction in DTS3 editing, and complete loss of DTS8 editing (Supplemental Fig. 4B).

### Assessing the genome-wide precision of NmeCas9 editing

All Cas9 orthologs described to date have some propensity to edit off-target sites lacking perfect complementarity to the programmed guide RNA, and considerable effort has been devoted to developing strategies (mostly with SpyCas9) to increase editing specificity (reviewed in [31, 34, 35]). In comparison with SpyCas9, orthologs such as NmeCas9 that employ longer guide sequences and that require longer PAMs have the potential for greater on-target specificity, possibly due in part to the lower density of near-cognate sequences. As an initial step in exploring this possibility, we used CRISPRseek [76] to perform a global analysis of potential NmeCas9 and SpyCas9 off-target sites with six or fewer mismatches in the human genome, using sgRNAs specific for DTS3, DTS7 and DTS8 (Fig. 5A) as representative queries. When allowing for permissive and semi-permissive PAMs (NGG, NGA, and NAG for SpyCas9; N_4_GHTT, N_4_GACT, N_4_GAYA, and N_4_GTCT for NmeCas9), potential off-target sites for NmeCas9 were predicted with two to three orders of magnitude lower frequency than for SpyCas9 (Table 2). Furthermore, NmeCas9 off-target sites with fewer than five mismatches were rare (two sites with four mismatches) for DTS7, and non-existent for DTS3 and DTS8 (Table 2). Even when we relaxed the NmeCas9 PAM requirement to N_4_GN_3_, which includes some PAMs that enable only background levels of targeting (e.g. N4GATC (Fig. 3A)), the vast majority of predicted off-target sites (>96%) for these three guides had five or more mismatches, and none had fewer than four mismatches (Fig. 5A). In contrast, the SpyCas9 guides targeting DTS3, DTS7, and DTS8 had 49, 54, and 62 predicted off-target sites with three or fewer mismatches, respectively (Table 2). As speculated previously [53, 54], these bioinformatic predictions suggest the intriguing possibility that the NmeCas9 genome editing system may induce very few undesired mutations, or perhaps none, even when targeting sites that induce substantial off-targeting with SpyCas9.

**Figure 5.**
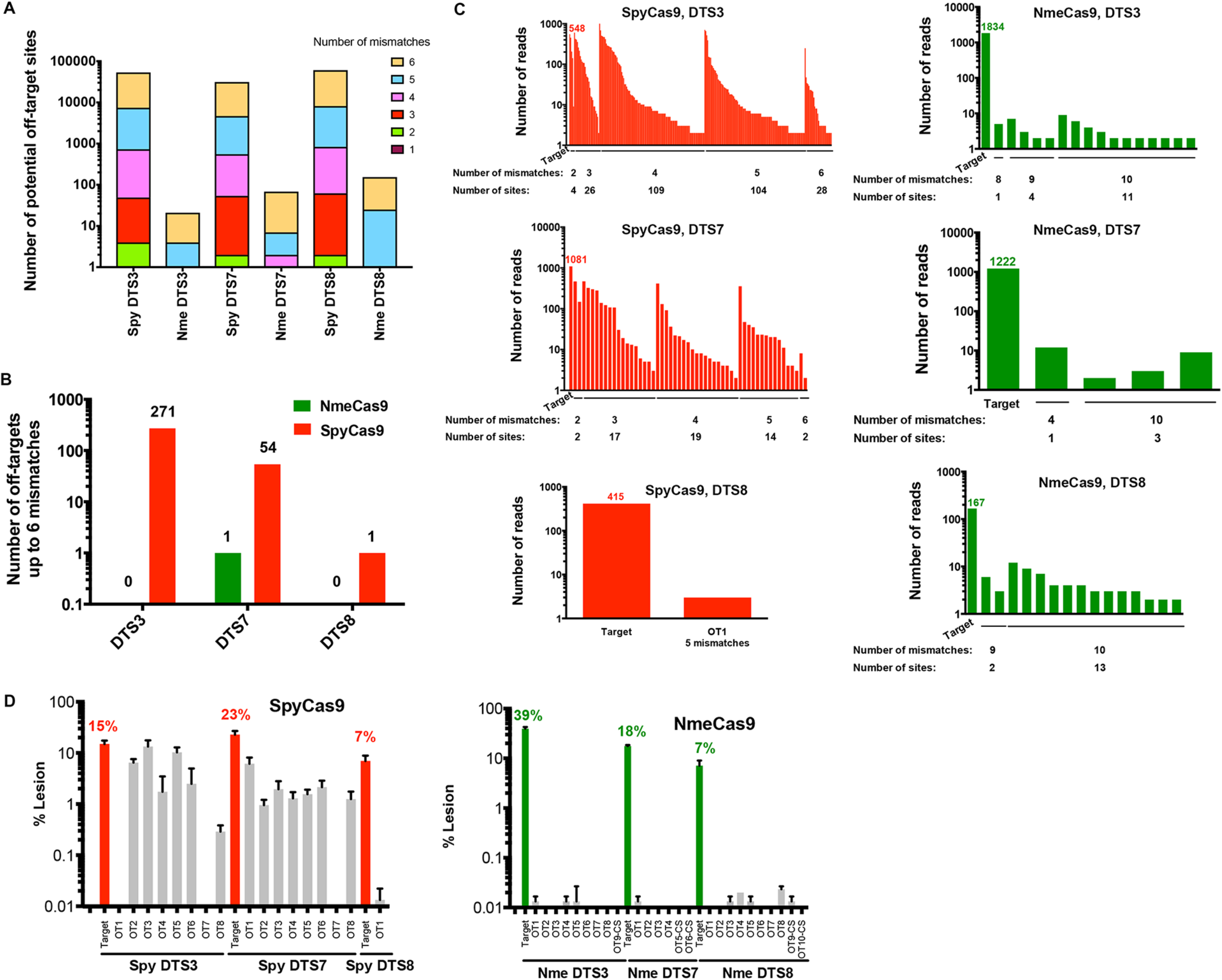
Bioinformatic and empirical comparison of NmeCas9 and SpyCas9 off-target sites within the human genome. (*A*) Genome-wide computational (CRISPRseek) predictions of off-target sites for NmeCas9 (with N4GN3 PAMs) and SpyCas9 (with NGG, NGA, and NAG PAMs) with DTS3, DTS7 and DTS8 sgRNAs. Predicted off-target sites were binned based on the number of mismatches (up to six) with the guide sequences. (*B*) GUIDE-Seq analysis of off-target sites in HEK293T cells with sgRNAs targeting DTS3, DTS7 and DTS8, using either SpyCas9 or NmeCas9, and with up to 6 mismatches to the sgRNAs. The numbers of detected off-target sites are indicated at the top of each bar. (*C*) Numbers of independent GUIDE-Seq reads for the on- and off-target sites for all six Cas9/sgRNA combinations from (*B*) (SpyCas9, red; NmeCas9, green), binned by the number of mismatches with the corresponding guide. (*D*) Targeted deep sequencing analysis of lesion efficiencies at on- and off-target sites from (*A*) or (*B*) with SpyCas9 (left, red) or NmeCas9 (right, green). Data for off-target sites are in grey. For SpyCas9, all off-target sites were chosen from (*B*) based on the highest GUIDE-Seq read counts for each guide (Supplemental Table 3). For NmeCas9, in addition to those candidate off-target sites obtained from GUIDE-Seq (*C*), we also assayed one or two potential off-target sites (designated with the “-CS” suffix) predicted by CRISPRseek as the closest near-cognate matches with permissive PAMs. Data are mean values ± s.e.m. from three biological replicates performed on different days.

**Table 2.** Number of predicted near-cognate sites in the human genome for the three dual target sites (DTS3, DTS7 and DTS8) analyzed in this study. These potential off-target sites differ from the on-target site by six or fewer mismatches, as listed on the left, and include the functional or semi-functional PAMs shown at the top.

| Number of mismatches | SpyCas9 sites<br>(NGG, NGA, NAG PAMs) |  |  | NmeCas9 sites (N<sub>4</sub>GATT, N<sub>4</sub>GCTT, N<sub>4</sub>GTTT, N<sub>4</sub>GACT, N<sub>4</sub>GATA, N<sub>4</sub>GTCT, N<sub>4</sub>GACA PAMs) |  |  |
| --- | --- | --- | --- | --- | --- | --- |
|  | DTS3 | DTS7 | DTS8 | DTS3 | DTS7 | DTS8 |
| 1 | 0 | 0 | 0 | 0 | 0 | 0 |
| 2 | 4 | 2 | 2 | 0 | 0 | 0 |
| 3 | 45 | 52 | 60 | 0 | 0 | 0 |
| 4 | 680 | 500 | 772 | 0 | 2 | 0 |
| 5 | 6,691 | 4,116 | 7,325 | 4 | 5 | 25 |
| 6 | 45,897 | 26,474 | 52,547 | 17 | 61 | 129 |
| **Total** | **53,317** | **31,144** | **60,706** | **21** | **68** | **154** |

Although bioinformatic predictions of off-targeting can be useful, it is well established that off-target profiles must be defined experimentally in a prediction-independent fashion due to our limited understanding of target specificity determinants, and the corresponding inability of algorithms to predict all possible sites successfully [31, 34, 35]. The need for empirical off-target profiling is especially acute with Cas9 orthologs that are far less thoroughly characterized than SpyCas9. A previous report used PCR amplification and high-throughput sequencing to detect the frequencies of lesions at 15-20 predicted NmeCas9 off-target sites for each of three guides in human cells, and found only background levels of indels in all cases, suggesting a very high degree of precision for NmeCas9 [54]. However, this report restricted its analysis to candidate sites with N4GNTT PAMs and three or fewer mismatches (or two mismatches combined with a 1-nt bulge) in the PAM-proximal 19 nucleotides, leaving open the possibility that legitimate off-target sites that did not fit these specific criteria remained unexamined. Accordingly, empirical and minimally-biased off-target profiles have never been generated for any NmeCas9/sgRNA combination, and the true off-target propensity of NmeCas9 therefore remains unknown. At the time we began this work, multiple methods for prediction-independent detection of off-target sites had been reported including GUIDE-seq, BLESS, Digenome-Seq, HTGTS, and IDLV capture, each with their own advantages and disadvantages (reviewed in [31, 34, 35]); additional methods (SITE-Seq [64], CIRCLE-seq [77], and BLISS [78]) have been reported more recently. Initially we chose to apply GUIDE-seq [63], which takes advantage of oligonucleotide incorporation into double-strand break sites, for defining the off-target profiles of both SpyCas9 and NmeCas9 when each is programmed to edit the DTS3, DTS7 and DTS8 sites (Fig. 4C-D) in the human genome.

After confirming that the co-transfected double-stranded oligodeoxynucleotide (dsODN) was incorporated efficiently at the DTS3, DTS7 and DTS8 sites during both NmeCas9 and SpyCas9 editing (Supplemental Fig. 4C), we then prepared GUIDE-seq libraries for each of the six editing conditions, as well as for the negative control conditions (i.e., in the absence of any sgRNA) for both Cas9 orthologs. The GUIDE-seq libraries were then subjected to high-throughput sequencing, mapped, and analyzed as described [79] (Fig. 5B-C). On-target editing with these guides was readily detected by this method, with the number of independent reads ranging from a low of 167 (NmeCas9, DTS8) to a high of 1,834 (NmeCas9, DTS3) (Fig. 5C and Supplemental Table 2).

For our initial analyses, we scored candidate sites as true off-targets if they yielded two or more independent reads and had six or fewer mismatches with the guide, with no constraints placed on the PAM match at that site. For SpyCas9, two of the sgRNAs (targeting DTS3 and DTS7) induced substantial numbers of off-target editing events (271 and 54 off-target sites, respectively (Fig. 5B)) under these criteria. The majority of these SpyCas9 off-target sites (88% and 77% for DTS3 and DTS7, respectively) were associated with a canonical NGG PAM. Reads were very abundant at many of these loci, and at five off-target sites (all with the DTS3 sgRNA) even exceeded the number of on-target reads (Fig. 5C). SpyCas9 was much more precise with the DTS8 sgRNA: we detected a single off-target site with five mismatches and an NGG PAM, and it was associated with only three independent reads, far lower than the 415 reads that we detected at the on-target site (Fig. 5C and Supplemental Table 2). Overall, the range of editing accuracies that we measured empirically for SpyCas9 — very high (e.g. DTS8), intermediate (e.g. DTS7), and poor (e.g. DTS3) – are consistent with the observations of other reports using distinct guides (reviewed in [31, 34, 35]).

In striking contrast, GUIDE-seq analyses with NmeCas9, programmed with sgRNAs targeting the exact same three sites, yielded off-target profiles that were exceptionally specific in all cases (Fig. 5B-C). For DTS3 and DTS8 we found no reads at any site with six or fewer guide mismatches; for DTS7 we found one off-target site with four mismatches (three of which were at the PAM-distal end; see Supplemental Table 2), and even at this site there were only 12 independent reads, ~100x fewer than the 1,222 reads detected at DTS7 itself. This off-target site was also associated with a PAM (N4GGCT) that would be expected to be poorly functional, though it could also be considered a “slipped” PAM with a more optimal consensus but variant spacing (N_5_GCTT). Purified, recombinant NmeCas9 has been observed to catalyze DNA cleavage *in vitro* at a site with a similarly slipped PAM [52]. To explore the off-targeting potential of NmeCas9 further, we decreased the stringency of our mapping to allow detection of off-target sites with up to 10 mismatches. Even in these conditions, only four (DTS7), 15 (DTS8), and 16 (DTS3) candidate sites were identified, most of which had only four or fewer reads (Fig. 5C) and were associated with poorly functional PAMs (Supplemental Table 2). We consider it likely that most if not all of these low-probability candidate off-target sites represent background noise caused by spurious priming and other sources of experimental error.

As an additional test of off-targeting potential, we repeated the DTS7 GUIDE-seq experiments with both SpyCas9 and NmeCas9, but this time using a different transfection reagent (Lipofectamine3000 rather than Polyfect). These repeat experiments revealed that >96% (29 out of 30) of off-target sites with up to five mismatches were detected under both transfection conditions for SpyCas9 (Supplemental Table 1). However, the NmeCas9 GUIDE-seq data showed no overlap between the potential sites identified under the two conditions, again suggesting that the few off-target reads that we did observe are unlikely to represent legitimate off-target editing sites.

To confirm the validity of the off-target sites defined by GUIDE-seq, we designed primers flanking candidate off-target sites identified by GUIDE-seq, PCR-amplified those loci following standard genome editing (i.e., in the absence of co-transfected GUIDE-seq dsODN) (3 biological replicates), and then subjected the PCR products to high-throughput sequencing to detect the frequencies of Cas9-induced indels. For this analysis we chose the top candidate off-target sites (as defined by GUIDE-seq read count) for each of the six cases (DTS3, DTS7 and DTS8, each edited by either SpyCas9 or NmeCas9). In addition, due to the low numbers of off-target sites and the low off-target read counts observed during the NmeCas9 GUIDE-seq experiments, we analyzed the top two predicted off-target sites for the three NmeCas9 sgRNAs, as identified by CRISPRseek (Fig. 5A and Table 2) [76]. On-target indel formation was detected in all cases, with editing efficiencies ranging from 7% (DTS8, with both SpyCas9 and NmeCas9) to 39% (DTS3 with NmeCas9) (Fig. 5D). At the off-target sites, our targeted deep-sequencing analyses largely confirmed our GUIDE-seq results: SpyCas9 readily induced indels at most of the tested off-target sites when paired with the DTS3 and DTS7 sgRNAs, and in some cases the off-target editing efficiencies approached those observed at the on-target sites (Fig. 5D). Although some SpyCas9 off-targeting could also be detected with the DTS8 sgRNA, the frequencies were much lower (<0.1% in all cases). Off-target lesions induced by NmeCas9 were far less frequent in all cases, even with the DTS3 sgRNA that was so efficient at on-target mutagenesis: many off-target sites exhibited lesion efficiencies that were indistinguishable from background, and never rose above ~0.02% (Fig. 5D). These results, in combination with the GUIDE-seq analyses described above, reveal wild-type NmeCas9 to be an exceptionally precise genome editing enzyme.

To explore NmeCas9 editing accuracy more deeply, we chose 16 additional NmeCas9 target sites across the genome, 10 with canonical N4GATT PAMs and six with variant functional PAMs (Supplemental Table 9). We then performed GUIDE-seq and analyses of NmeCas9 editing at these sites. GUIDE-seq analysis readily revealed editing at each of these sites, with on-target read counts ranging from ~ 100 to ~5,000 reads (Fig. 6A). More notably, off-target reads were undetectable by GUIDE-seq with 14 out of the 16 sgRNAs (Fig. 6B). Targeted deep sequencing of PCR amplicons, which is a more quantitative readout of editing efficiency than either GUIDE-seq or T7E1 analysis, confirmed on-target editing in all cases, with indel efficiencies ranging from ~5-85% (Fig. 6C).

**Figure 6.**
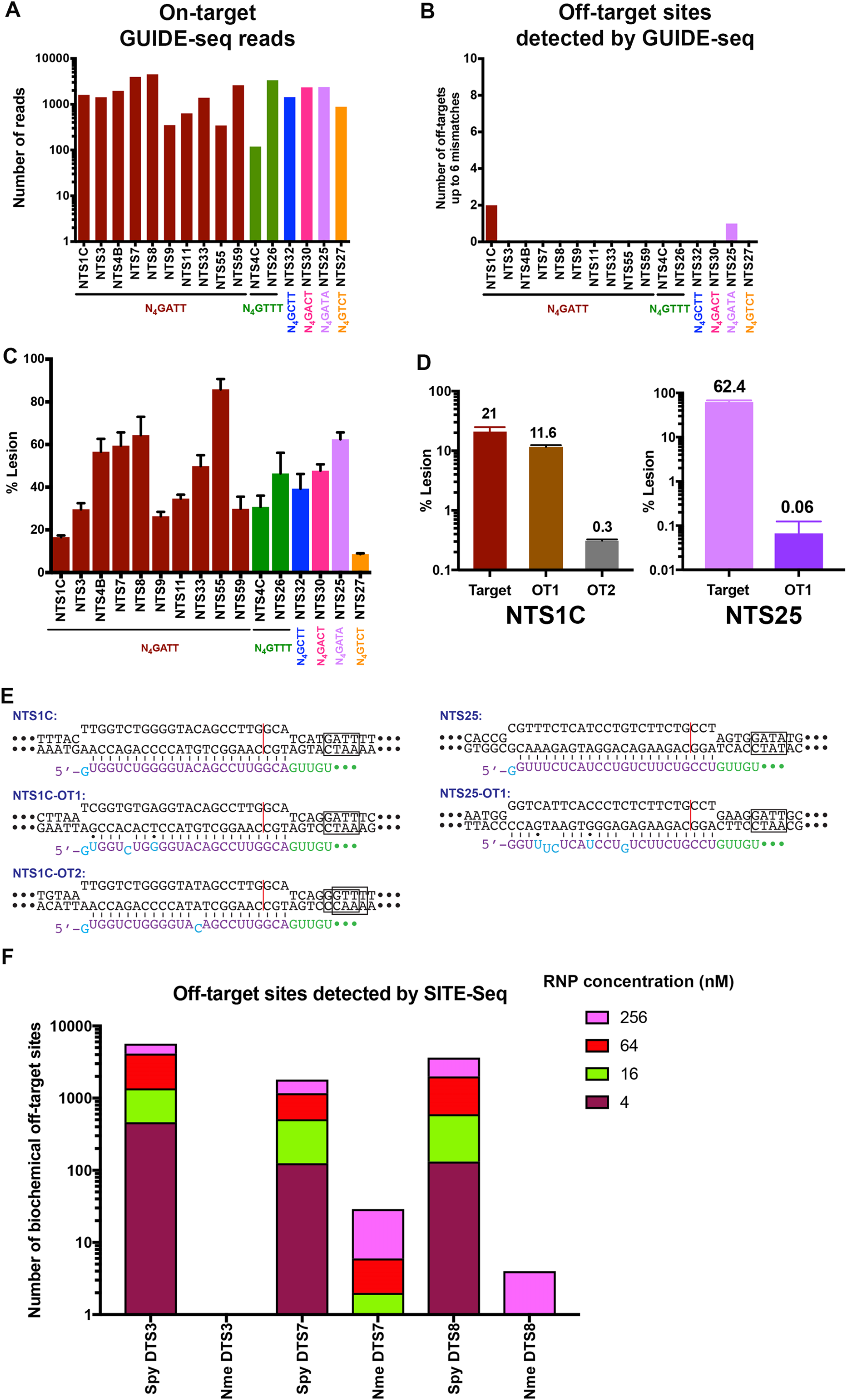
Off-target analyses for additional NmeCas9 sgRNAs, targeting sites with consensus and variant PAMs. (*A*) Number of GUIDE-Seq reads for the on-target sites, with the PAM sequences for each site indicated underneath. (*B*) Number of GUIDE-Seq-detected off-target sites using the Bioconductor package GUIDEseq version 1.1.17 [79] with default settings except that PAM.size = 8, PAM =“NNNNGATT”, min.reads = 2, max.mismatch = 6, allowed.mismatch.PAM = 4, PAM.pattern =“NNNNNNNN$”, BSgenomeName = Hsapiens, txdb = TxDb.Hsapiens.UCSC.hg19.knownGene, orgAnn = org.Hs.egSYMBOL gRNA.size was set to length of the gRNA used, and various number of 0’s were added at the beginning of weights to make the length of weights equal to the gRNA size. For example, for gRNA with length 24, weights = c(0,0,0,0,0, 0, 0.014, 0, 0, 0.395, 0.317, 0, 0.389, 0.079, 0.445, 0.508, 0.613, 0.851, 0.732, 0.828, 0.615, 0.804, 0.685, 0.583) for all sixteen sgRNAs used in (*A*). (*C*) Lesion efficiencies for the on-target sites as measured by PCR and high-throughput sequencing. Data are mean values ± s.e.m. from three biological replicates performed on different days. (*D*) NmeCas9 lesion efficiencies at the NTS 1C (left) and NTS25 (right) on-target sites, and at the off-target sites detected by GUIDE-Seq from (*B*), as measured by PCR and high-throughput sequencing. Data are mean values ± s.e.m. from three biological replicates performed on different days. (*E*) Schematic diagrams of NmeCas9 sgRNA/DNA R-loops for the NTS1C (left) and NTS25 (right) sgRNAs, at the GUIDE-Seq-detected on- and off-target sites. Black, DNA residues; boxed nts, PAM; red line, NmeCas9 cleavage site; cyan and purple, mismatch/wobble and complementary nts (respectively) in the NmeCas9 sgRNA guide region; green, NmeCas9 sgRNA repeat nts. (*F*)Comparison of NmeCas9 and SpyCas9 biochemical off-target sites using SITE-Seq analysis.

The two guides with off-target activity (NTS1C and NTS25) had only two and one off-target sites, respectively (Fig. 6B and Supplemental Fig. 5). Off-target editing was confirmed by high-throughput sequencing and analysis of indels (Fig. 6D). Compared with the on-target site (perfectly matched at all positions other than the 5’-terminal guide nucleotide, and with an optimal N_4_GATT PAM), the efficiently targeted NTS1C-OT1 had two wobble pairs and one mismatch (all in the nine PAM-distal nucleotides), as well as a canonical N_4_GATT PAM (Fig. 6E and Supplemental Table 2). The weakly edited NTS1C-OT2 site had only a single mismatch (at the 11^th^ nucleotide, counting in the PAM-distal direction), but was associated with a non-canonical N4GGTT (or a “slipped” N5GTTT) PAM (Fig. 6E and Supplemental Table 2). NTS25 with an N_4_GATA PAM was the other guide with a single off-target site (NTS25-OT1), where NmeCas9 cleaved and edited up to ~1,000x less efficiently than at the on-target site (Fig. 6D). This minimal amount of off-target editing arose despite the association of NTS25-OT1 with an optimal N4GATT PAM, unlike the variant N4GATA PAM that flanks the on-target site. Overall, our GUIDE-seq and sequencing-based analyses demonstrate that NmeCas9 genome editing is exceptionally accurate: we detected and confirmed cellular off-target editing with only two of the 19 guides tested, and even in those two cases, only one or two off-target sites could be found for each. Furthermore, of the three bona fide off-target sites that we identified, only one generated indels at substantial frequency (11.6%); indel frequencies were very modest (0.3% or lower) at the other two off-target sites.

We next sought to corroborate and expand on our GUIDE-seq results with a second prediction-independent method. We applied the SITE-Seq^™^ (Caribou Biosciences, Inc., Berkeley, CA) assay, a biochemical-based method that does not rely on cellular events such as DNA repair, thus potentially enabling a more thorough profiling of genome-wide specificity [63]. SITE-Seq libraries were prepared for the three dual target sites with both Cas9 orthologues as well as for twelve of the NmeCas9-only target sites. SITE-Seq was performed on HEK293T genomic DNA (gDNA) treated with a range of RNP concentrations (4 nM – 256 nM) previously shown to discriminate high and low probability cellular off-targets [63]. Finally, the resulting libraries were sequenced, aligned, and then analyzed as previously described [63].

Negative controls without RNP recovered zero sites across any concentrations, whereas SpyCas9 assembled with sgRNAs targeting DTS3, DTS7, or DTS8 recovered hundreds (at 4 nM RNP) to thousands (at 256 nM RNP) of biochemical off-target sites (Fig. 6F). In contrast, NmeCas9 assembled with sgRNAs targeting the same three sites recovered only their on-target sites at 4 nM RNP and at most 29 off-target sites at 256 nM RNP (Fig. 6F). Moreover, the 12 additional NmeCas9 target sites showed similarly high specificity: eight samples recovered only the on-target sites at 4 nM RNP and six of those recovered no more than nine off-targets at 256 nM RNP (Supplemental Fig. 6A). Across NmeCas9 RNPs, off-target sequence mismatches appeared enriched in the 5’ end of the sgRNA target sequence (Supplemental Table 4). Finally, three of the NmeCas9 RNPs (NTS30, NTS4C, and NTS59) required elevated concentrations to retrieve their on-targets, potentially due to poor sgRNA transcription and/or RNP assembly. These RNPs were therefore excluded from further analysis.

We next performed cell-based validation experiments to investigate whether any of the biochemical off-targets were edited in cells. Since NmeCas9 recovered only ~ 100 biochemical off-targets across all RNPs and concentrations, we could examine each site for editing in cells. SpyCas9 generated >10,000 biochemical off-targets across all DTS samples, preventing comprehensive cellular profiling. Therefore, for each RNP we selected 96 of the high cleavage sensitivity SITE-Seq sites (i.e., recovered at all concentrations tested in SITE-Seq) for examination, as we predicted those were more likely to accumulate edits in cells [63] (Supplemental Table 5). Sites were randomly selected within this cohort and only included a subset of the GUIDE-seq validation test set sites (1/8 and 5/8 overlapping sites for DTS3 and DTS7, respectively). Additionally, SITE-Seq and GUIDE-seq validations were performed on the same gDNA samples to facilitate comparisons between data sets.

Across all NmeCas9 RNPs, only three cellular off-targets were observed. These three all belonged to the NTS1C RNP, and two of them had also been detected with GUIDE-seq. Of note, all high cleavage sensitivity SITE-Seq sites (i.e., all on-targets and the single prominent NTS1C off-target, NTS1C-OT1) showed editing in cells. Conversely, SITE-Seq sites with low cleavage sensitivity, defined as being recovered at only 64 nM and/or 256 nM RNP, were rarely found as edited (2/93 sites). Importantly, this suggests that we identified all or the clear majority of NmeCas9 cellular off-targets, albeit at our limit of detection. Across all SpyCas9 RNPs, 14 cellular off-targets were observed (8/70 sites for DTS3, 6/83 sites for DTS7, and 0/79 sites for DTS8) (Supplemental Table 5). Since our data set was only a subset of the total number of high cleavage sensitivity SITE-Seq sites, and excluded many of the GUIDE-seq validated sites, we expect that sequencing all SITE-Seq sites would uncover additional cellular off-targets. Taken together, these data corroborate our GUIDE-seq results, suggesting that NmeCas9 can serve as a highly specific genome editing platform.

### Indel spectrum at NmeCas9-edited sites

Our targeted deep sequencing data at the three dual target sites (Fig. 5D, Supplemental Fig. 4A and Supplemental Table 5) enabled us to analyze the spectrum of insertions and deletions generated by NmeCas9, in comparison with those of SpyCas9 when editing the exact same sites (Supplemental Figs. 6B and 7-9). Although small deletions predominated at all three sites with both Cas9 orthologs, the frequency of insertions was lower for NmeCas9 than it was with SpyCas9 (Supplemental Figs. 6B and 7-9). For both SpyCas9 and NmeCas9, the vast majority of insertions were only a single nucleotide (Supplemental Fig. 8). The sizes of the deletions varied from one target site to the other for both Cas9 orthologs. Our data suggest that at Cas9 edits, deletions predominated over insertions and the indel size varies considerably site to site (Supplemental Figs. 6B, 10 and 11).

### Truncated sgRNAs reduce off-target cleavage by NmeCas9

Although NmeCas9 exhibits very little propensity to edit off-target sites, for therapeutic applications it may be desirable to suppress even the small amount of off-targeting that occurs (Fig. 6). Several strategies have been developed to suppress off-targeting by SpyCas9 [31, 34, 35], some of which could be readily applied to other orthologs. For example, truncated sgRNAs (tru-sgRNAs) sometimes suppress off-target SpyCas9 editing more than they suppress on-target editing [69]. Because 5’-terminal truncations are compatible with NmeCas9 function (Fig. 2), we tested whether NmeCas9 tru-sgRNAs can have similar suppressive effects on off-target editing without sacrificing on-target editing efficiency.

First, we tested whether guide truncation can lead to NmeCas9 editing at novel off-target sites (i.e. at off-target sites not edited by full-length guides), as reported previously for SpyCas9 [69]. Our earlier tests of NmeCas9 on-target editing with tru-sgRNAs used guides targeting the NTS33 (Fig. 2B-C) and NTS32 (Fig. 2D-E) sites. GUIDE-seq did not detect any NmeCas9 off-target sites during editing with full-length NTS32 and NTS33 sgRNAs (Fig. 6). We again used GUIDE-seq with a subset of the validated NTS32 and NTS33 tru-sgRNAs to determine whether NmeCas9 guide truncation leads to off-target editing at new sites, and found none (Supplemental Fig. 12). Although we cannot rule out the possibility that other NmeCas9 guides could be identified that yield novel off-target events upon truncation, our results suggest that *de novo* off-targeting by NmeCas9 tru-sgRNAs is unlikely to be a pervasive problem.

The most efficiently edited off-target site from our previous analyses was NTS1C-OT1, providing us with our most stringent test of off-target suppression. When targeted by the NTS1C sgRNA, NTS1C-OT1 has one rG-dT wobble pair at position −16 (i.e., at the 16^th^ base pair from the PAM-proximal end of the R-loop), one rC-dC mismatch at position -19, and one rU-dG wobble pair at position −23 (Fig. 6E). We generated a series of NTS1C-targeting sgRNAs with a single 5’-terminal G (for U6 promoter transcription) and spacer complementarities ranging from 24 to 15 nucleotides (GN_24_ to GN_15_, Supplemental Fig. 13A, top panel). Conversely, we designed a similar series of sgRNAs with perfect complementarity to NTS1C-OT1 (Supplemental Fig. 13B, top panel). Consistent with our earlier results with other target sites (Fig. 2), T7E1 analyses revealed that both sets of guides enabled editing of the perfectly-matched on-target site with truncations down to 19 nucleotides (GN18), but that shorter guides were inactive. On-target editing efficiencies at both sites were comparable across the seven active guide lengths (GN24 through GN18), with the exception of slightly lower efficiencies with the GN19 guides (Supplemental Fig. 13A & B, middle and bottom panels).

We then used targeted deep sequencing to test whether off-target editing is reduced with the truncated sgRNAs. With both sets of sgRNAs (perfectly complementary to either NTS 1C or NTS1C-OT1), we found that off-targeting at the corresponding near-cognate site persisted with the four longest guides (GN24, GN23, GN22, GN21; Fig. 7). However, off-targeting was abolished with the GN20 guide, without any significant reduction in on-target editing efficiencies (Fig. 7). Off-targeting was also absent with the GN19 guide, though on-target editing efficiency was compromised. These results, albeit from a limited data set, indicate that truncated sgRNAs (especially those with 20 or 19 base pairs of guide/target complementarity, 4-5 base pairs fewer than the natural length) can suppress even the limited degree of off-targeting that occurs with NmeCas9.

**Figure 7.**
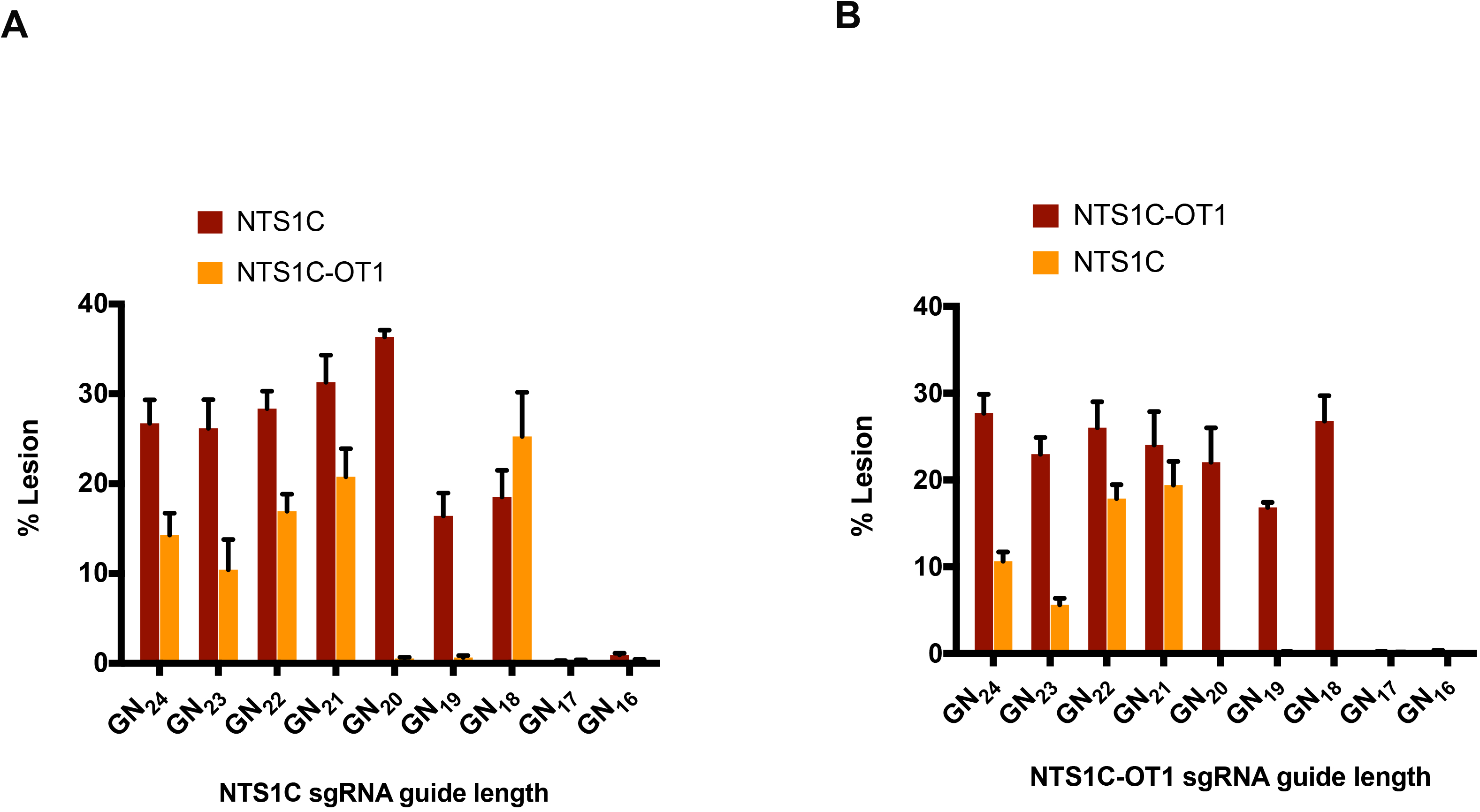
Guide truncation can suppress off-target editing by NmeCas9. (*A*) Lesion efficiencies at the NTS1C (on-target, red) and NTS1C-OT1 (off-target, orange) genomic sites, after editing by NmeCas9 and NTS1C sgRNAs of varying lengths, as measured by PCR and high-throughput sequencing. Data are mean values ± s.e.m. from three biological replicates performed on different days. (*B*) As in (*A*), but using sgRNAs perfectly complementary to the NTS1C-OT1 genomic site.

Unexpectedly, even though off-targeting at NTS1C-OT1 was abolished with the GN20 and GN19 truncated NTS 1C sgRNAs, truncating by an additional nucleotide (to generate the GN18 sgRNA) once again yielded NTS1C-OT1 edits (Fig. 7A). This could be explained by the extra G residue at the 5’-terminus of each sgRNA in the truncation series (Supplemental Fig. 13). With the NTS 1C GNı_9_ sgRNA, both the 5’-terminal G residue and the adjacent C residue are mismatched with the NTS1C-OT1 site. In contrast, with the GN_18_ sgRNA, the 5’-terminal G is complementary to the off-target site. In other words, with the NTS1C GN_19_ and GN_18_ sgRNAs, the NTS1C-OT1 off-target interactions (which are identical in the PAM-proximal 17 nucleotides) include two additional nucleotides of non-complementarity or one additional nucleotide of complementarity, respectively. Thus, the more extensively truncated GN18 sgRNA has *greater* complementarity with the NTS 1C-OT1 site than the GN_19_ sgRNA, explaining the reemergence of off-target editing with the former. This observation highlights the fact that the inclusion of a 5’-terminal G residue that is mismatched with the on-target site, but that is complementary to a C residue at an off-target site, can limit the effectiveness of a truncated guide at suppressing off-target editing, necessitating care in truncated sgRNA design when the sgRNA is generated by cellular transcription. This issue is not a concern with sgRNAs that are generated by other means (e.g. chemical synthesis) that do not require a 5’-terminal G. Overall, our results demonstrate that NmeCas9 genome editing is exceptionally precise, and even when rare off-target editing events occur, tru-sgRNAs can provide a simple and effective way to suppress them.

## DISCUSSION

The ability to use Type II and Type V CRISPR-Cas systems as RNA-programmable DNA-cleaving systems [13, 14, 27] is revolutionizing many aspects of the life sciences, and holds similar promise for biotechnological, agricultural, and clinical applications. Most applications reported thus far have used a single Cas9 ortholog (SpyCas9). Thousands of additional Cas9 orthologs have also been identified [28], but only a few have been characterized, validated for genome engineering applications, or both. Adding additional orthologs promises to increase the number of targetable sites (through new PAM specificities), extend multiplexing possibilities (for pairwise combinations of Cas9 orthologs with orthogonal guides), and improve deliverability (for the more compact Cas9 orthologs). In addition, some Cas9s may show mechanistic distinctions (such as staggered vs. blunt dsDNA breaks) [80], greater protein stability *in vivo*, improved control mechanisms (e.g. via multiple anti-CRISPRs that act at various stages of the DNA cleavage pathway) [57, 58, 60, 61, 81-83], and other enhancements. Finally, some may exhibit a greater natural propensity to distinguish between on- vs. off-target sites during genome editing applications, obviating the need for extensive engineering (as was necessary with SpyCas9) to attain the accuracy needed for many applications, especially therapeutic development.

Here we have further defined the properties of NmeCas9 during editing in human cells, including validation and extension of previous analyses of guide length and PAM requirements [46, 53, 54]. Intriguingly, the tolerance to deviations from the N_4_G(A/C)TT natural PAM consensus [51] observed *in vitro* and in bacterial cells [46, 52] is considerably reduced in the mammalian context, i.e. fewer PAM variations are permitted during mammalian editing. The basis for this context-dependent difference is not clear but may be due in part to the ability to access targets within eukaryotic chromatin, or to decreased expression levels relative to potential DNA substrates, since lower SpyCas9/sgRNA concentrations have been shown to improve accuracy [30, 84, 85]. We have also found that steady-state NmeCas9 levels in human cells are markedly increased in the presence of its cognate sgRNA, suggesting that sgRNA-loaded NmeCas9 is more stable than *apo* NmeCas9. An increased proteolytic sensitivity of *apo* Cas9 relative to the sgRNA-bound form has been noted previously for a different Type II-C ortholog *(Corynebacterium diphtheria*Cas9, CdiCas9 [55]).

A previous report indicated that NmeCas9 has high intrinsic accuracy, based on analyses of candidate off-target sites that were predicted bioinformatically [54]. However, the true genome-wide accuracy of NmeCas9 was not assessed empirically, as is necessary given well-established imperfections in bioinformatic predictions of off-targeting [31, 34, 35]. We have used GUIDE-seq [63] and SITE-Seq [64] to define the genome-wide accuracy of wild-type NmeCas9, including side-by-side comparisons with wildtype SpyCas9 during editing of identical on-target sites. We find that NmeCas9 is a consistently high-accuracy genome editor, with off-target editing undetectable above background with 17 out of 19 analyzed sgRNAs, and only one or three verified off-target edits with the remaining two guides. We observed this exquisite specificity by NmeCas9 even with sgRNAs that target sites (DTS3 and DTS7 (see Fig. 5)) that are highly prone to off-target editing when targeted with SpyCas9. Of the four off-target sites that we validated, three accumulated < 1% indels. Even with the one sgRNA that yielded a significant frequency of off-target editing (NTS1C, which induced indels at NTS1C-OT1 with approximately half the efficiency of on-target editing), the off-targeting with wild-type NmeCas9 could be easily suppressed with truncated sgRNAs. Our ability to detect NTS25-OT1 editing with GUIDE-seq, despite its very low (0.06%) editing efficiency based on high-throughput sequencing, indicates that our GUIDE-seq experiments can identify even very low-efficiency off-target editing sites. Similar considerations apply to our SITE-Seq analyses. We observed high accuracy even when NmeCas9 is delivered by plasmid transfection, a delivery method that is associated with higher off-target editing than more transient delivery modes such as RNP delivery [86, 87].

The two Type II-C Cas9 orthologs (NmeCas9 and CjeCas9) that have been validated for mammalian genome editing and assessed for genome-wide specificity [47, 54] (this work) have both proven to be naturally hyper-accurate. Both use longer guide sequences than the 20-nucleotide guides employed by SpyCas9, and both also have longer and more restrictive PAM requirements. For both Type II-C orthologs, it is not yet known whether the longer PAMs, longer guides, or both account for the limited off-target editing. Type II-C Cas9 orthologs generally cleave dsDNA more slowly than SpyCas9 [49, 55], and it has been noted that lowering *k*_cat_ can, in some circumstances, enhance specificity [88]. Whatever the mechanistic basis for the high intrinsic accuracy, it is noteworthy that it is a property of the native proteins, without a requirement for extensive engineering. This adds to the motivation to identify more Cas9 orthologs with human genome editing activity, as it suggests that it may be unnecessary in many cases (perhaps especially among Type II-C enzymes) to invest heavily in structural and mechanistic analyses and engineering efforts to attain sufficient accuracy for many applications and with many desired guides, as was done with (for example) SpyCas9 [32, 33, 37, 38, 65]. Although Cas9 orthologs with more restrictive PAM requirements (such as NmeCas9, CjeCas9, and GeoCas9) by definition will afford lower densities of potential target sites than SpyCas9 (which also usually affords the highest on-target editing efficiencies among established Cas9 orthologs), the combined targeting possibilities for multiple such Cas9s will increase the targeting options available within a desired sequence window, with little propensity for off-targeting. The continued exploration of natural Cas9 variation, especially for those orthologs with other advantages such as small size and anti-CRISPR off-switch control, therefore has great potential to advance the CRISPR genome editing revolution.

## CONCLUSIONS

NmeCas9 is an intrinsically high-accuracy genome editing enzyme in mammalian cells, and the limited off-target editing that occurs can (at least in some cases) be suppressed by guide truncation. Continued exploration of Cas9 orthologs could therefore yield additional enzymes that do not require extensive characterization and engineering to prevent off-target editing.

## METHODS

### Plasmids

Two plasmids for the expression of NmeCas9 were used in this study. The first construct (used in Figs. 1 and 2) was derived from the plasmid pSimpleII where NmeCas9 was cloned under the control of the elongation factor-1α promoter, as described previously [53]. The *Cas9* gene in this construct expresses a protein with two NLSs and an HA tag. To make an all-in-one expression plasmid, a fragment containing a *BsmB*I-crRNA cassette linked to the tracrRNA by six nucleotides, under the control of U6 RNA polymerase III promoter, was synthesized as a gene block (Integrated DNA Technologies) and inserted into pSimpleII, generating the pSimpleII-Cas9-sgRNA-*BsmB*I plasmid that includes all elements needed for editing. To insert specific spacer sequence into the crRNA cassette, synthetic oligonucleotides were annealed to generate a duplex with overhangs compatible with those generated by *BsmB*I digestion of the pSimpleII-Cas9-sgRNA-*BsmB*I plasmid. The insert was then ligated into the *BsmB*I-digested plasmid. For Figs. 3-7, NmeCas9 and SpyCas9 constructs were expressed from the pCS2-Dest Gateway plasmid under the control of the CMV IE94 promoter [89]. All sgRNAs used with pCS2-Dest-Cas9 were driven by the U6 promoter in pLKO.1-puro [90]. The M427 GFP reporter plasmid [66] was used as described [65]. Cell culture, transfection, and transduction HEK293T were cultured in DMEM with 10% FBS and 1% Penicillin/Streptomycin (Gibco) in a 37°C incubator with 5% CO2. K562 cells were grown in the same conditions but using IMDM. HFF cells were grown in the same conditions but using DMEM with Glutamax and 20% FBS without antibiotics. mESCs were grown in DMEM supplemented with 10% FBS, glutamine beta-ME and LIF. For transient transfection, we used early to mid-passage cells (passage number 4-18). Approximately 1.5 × 10^5^ cells were transfected with 150 ng Cas9-expressing plasmid, 150 ng sgRNA-expressing plasmid and 10 ng mCherry plasmid using Polyfect transfection reagent (Qiagen) in a 24-well plate according to the manufacturer’s protocol. For the GFP reporter assay, 100 ng M427 plasmid was included in the co-transfection mix. Transduction was done as described previously [91].

### Western blotting

48 h after transfection, cells were harvested and lysed with 50 μl of RIPA buffer. Protein concentration was determined with the BCA kit (Thermo Scientific) and 12 μg of proteins were used for electrophoresis and blotting. The blots were probed with anti-HA (Sigma, H3663) and anti-GAPDH (Abcam, ab9485) as primary antibodies, and then with horseradish peroxidase–conjugated anti-mouse IgG (Thermoscientific, 62-6520) or anti-rabbit IgG (Biorad, 1706515) secondary antibodies, respectively. Blots were visualized using the Clarity Western ECL substrate (Biorad, 170-5060).

### Flow cytometry

The GFP reporter was used as described previously [65]. Briefly, cells were harvested 48 hours after transfection and used for FACS analysis (BD Accuri 6C). To minimize the effects of differences in the efficiency of transfection among samples, cells were initially gated for mCherry-expression, and the percentage of GFP-expressing cells were quantified within mCherry positive cells. All experiments were performed in triplicate with data reported as mean values with error bars indicating the standard error of the mean (s.e.m.).

### Genome editing

72 hours after transfection, genomic DNA was extracted via the DNeasy Blood and Tissue kit (Qiagen), according to the manufacturer’s protocol. 50 ng DNA was used for PCR-amplification using primers specific for each genomic site (Supplemental Table 9) with High Fidelity 2X PCR Master Mix (New England Biolabs). For T7E1 analysis, 10 μl of PCR product was hybridized and treated with 0.5 μl T7 Endonuclease I (10 U/μl, New England Biolabs) in 1X NEB Buffer 2 for 1 hour. Samples were run on a 2.5% agarose gel, stained with SYBR-safe (ThermoFisher Scientific), and quantified using the ImageMaster-TotalLab program. Indel percentages are calculated as previously described [92, 93]. Experiments for T7E1 analysis are performed in triplicate with data reported as mean ± s.e.m. For indel analysis by TIDE, 20 ng of PCR product is purified and then sequenced by Sanger sequencing. The trace files were subjected to analysis using the TIDE web tool (https://tide.deskgen.com).

### Expression and purification of NmeCas9

NmeCas9 was cloned into the pMCSG7 vector containing a T7 promoter followed by a 6xHis tag and a tobacco etch virus (TEV) protease cleavage site. Two NLSs on the C-terminus of NmeCas9 and another NLS on the N-terminus were also incorporated. This construct was transformed into the Rosetta 2 DE3 strain of *E. coli*. Expression of NmeCas9 was performed as previously described for SpyCas9 [14]. Briefly, a bacterial culture was grown at 37°C until an OD600 of 0.6 was reached. At this point the temperature was lowered to 18°C followed by addition of 1 mM Isopropyl β-D-1-thiogalactopyranoside (IPTG) to induce protein expression. Cells were grown overnight, and then harvested for purification. Purification of NmeCas9 was performed in three steps: Nickel affinity chromatography, cation exchange chromatography, and size exclusion chromatography. The detailed protocols for these can be found in [14].

### RNP delivery of NmeCas9

RNP delivery of NmeCas9 was performed using the Neon transfection system (ThermoFisher). Approximately 40 picomoles of NmeCas9 and 50 picomoles of sgRNA were mixed in buffer R and incubated at room temperature for 30 minutes. This preassembled complex was then mixed with 50,000 – 150,000 cells, and electroporated using 10 μL Neon tips. After electroporation, cells were plated in prewarmed 24-well plates containing the appropriate culture media without antibiotics. The number of cells used and pulse parameters of electroporation were different for different cell types tested. The number of cells used were 50,000, 100,000, and 150,000 for PLB985 cells, HEK293T cells, and K562/HFF cells respectively. Electroporation parameters (voltage, width, number of pulses) were 1150 v, 20 ms, 2 pulses for HEK293T cells; 1000 v, 50 ms, 1 pulse for K562 cells; 1350 v, 35 ms, 1 pulse for PLB985 cells; and 1700 v, 20 ms, 1pulse for HFF cells.

### γH2AX immunofluorescence staining and flow cytometry

For immunofluorescence, mouse embryonic stem cells (mESCs) were crosslinked with 4% paraformaldehyde and stained with anti-γH2AX (LP BIO, AR-0149-200) as primary antibody and Alexa Fluor^®^ 488 goat anti-rabbit IgG (Invitrogen, A11034) as secondary antibody. DNA was stained with DAPI. For a positive control, E14 cells were irradiated with 254 nm UV light (3 mJ/cm2). Images were taken by a Nikon Eclipse E400 and representative examples were chosen.

For flow cytometry, cells were fixed with 70% ethanol, primary and secondary antibody were as described above for immunofluorescence, and DNA was stained with propidium iodide. Cells were analyzed by BD FACSCalibur. The box plot was presented with the bottom line of the box representing the first quartile, the band inside box indicating the median, the top line being the third quartile, the bottom end of whisker denoting data of first quartile minus 1.5 times of interquartile range (no less than 0), and the top end of the whisker indicating data of third quartile plus 1.5 times of interquartile. Outliers are not shown. All experiments were performed in duplicate.

### CRISPRseek analysis of potential off-target sites

Global off-target analyses for DTS3, DTS7, and DTS8 with NmeCas9 sgRNAs were performed using the Bioconductor package CRISPRseek 1.9.1 [76] with parameter settings tailored for NmeCas9. Specifically, all parameters are set as default except the following: gRNA.size = 24, PAM =“NNNNGATT”, PAM.size =8, RNA.PAM.pattern =“NNNNGNNN$”, weights = c(0, 0, 0, 0, 0, 0, 0.014, 0, 0, 0.395, 0.317, 0, 0.389, 0.079, 0.445, 0.508, 0.613, 0.851, 0.732, 0.828, 0.615, 0.804, 0.685, 0.583), max.mismatch =6, allowed.mismatch.PAM = 7, topN = 10000, min.score = 0. This setting means that all seven permissive PAM sequences (N4GATT, N4GCTT, N4GTTT, N4GACA, N4GACT, N4GATA, N4GTCT) were allowed and all off-targets with up to 6 mismatches were collected [the sgRNA length was changed from 20 to 24; four additional zeros were added to the beginning of the weights series to be consistent with the gRNA length of 24; and topN (the number of off-target sites displayed) and min.score (the minimum score of an off-target to be included in the output) were modified to enable identification of all off-target sites with up to 6 mismatches]. Predicted off-target sites for DTS3, DTS7, and DTS8 with SpyCas9 sgRNAs were obtained using CRISPRseek 1.9.1 default settings for SpyCas9 (with NGG, NAG, and NGA PAMs allowed). Batch scripts for high-performance computing running the IBM LSF scheduling software are included in the supplemental section. Off-target sites were binned according to the number of mismatches relative to the on-target sequence. The numbers of off-targets for each sgRNA were counted and plotted as pie charts.

### GUIDE-seq

We performed GUIDE-seq experiment with some modifications to the original protocol [63], as described [65]. Briefly, in 24-well format, HEK293T cells were transfected with 150 ng of Cas9, 150 ng of sgRNA, and 7.5 pmol of annealed GUIDE-seq oligonucleotide using Polyfect transfection reagent (Qiagen) for all six guides (DTS3, DTS7 and DTS8 for both the NmeCas9 and SpyCas9 systems). Experiments with DTS7 sgRNAs were repeated using Lipofectamine 3000 transfection reagent (Invitrogen) according to the manufacturer’s protocol. 48 h after transfection, genomic DNA was extracted with a DNeasy Blood and Tissue kit (Qiagen) according to the manufacturer protocol. Library preparation, sequencing, and read analyses were done according to protocols described previously [63, 65]. Only sites that harbored a sequence with up to six or ten mismatches with the target site (for SpyCas9 or NmeCas9, respectively) were considered potential off-target sites. Data were analyzed using the Bioconductor package GUIDEseq version 1.1.17 (Zhu et al., 2017). For SpyCas9, default setting was used except that min.reads = 2, max.mismatch = 6, allowed.mismatch.PAM = 2, PAM.pattern =“NNN$”, BSgenomeName = Hsapiens, txdb = TxDb.Hsapiens.UCSC.hg19.knownGene, orgAnn = org.Hs.egSYMBOL For NmeCas9, default setting was used except that PAM.size = 8, PAM =“NNNNGATT”, min.reads = 2, allowed.mismatch.PAM = 4, PAM.pattern =“NNNNNNNN$”, BSgenomeName = Hsapiens, txdb = TxDb.Hsapiens.UCSC.hg19.knownGene, orgAnn = org.Hs.egSYMBOL. NmeCas9 dataset was analyzed twice with max.mismatch = 6 and max.mismatch = 10 respectively. The gRNA.size was set to the length of the gRNA used, and various number of 0’s was added at the beginning of weights to make the length of weights equal to the gRNA size. For example, for gRNA with length 24, weights = c(0,0,0,0,0, 0, 0.014, 0, 0, 0.395, 0.317, 0, 0.389, 0.079, 0.445, 0.508, 0.613, 0.851, 0.732, 0.828, 0.615, 0.804, 0.685, 0.583) (Zhu et al., 2017). These regions are reported in Supplemental Table 2.

### SITE-Seq

We performed the SITE-Seq assay as described previously [63]. In 50 mL conical tubes, high molecular weight genomic DNA (gDNA) was extracted from HEK293T cells using the Blood and Cell Culture DNA Maxi Kit (Qiagen) according to the manufacturer’s protocol. sgRNAs for both NmeCas9 and SpyCas9 RNP assembly were transcribed from PCR-assembled DNA templates containing T7 promoters. Oligo sequences used in DNA template assembly can found be in Supplemental Table 8. PCR reactions were performed using Q5 Hot Start High-Fidelity 2X Master Mix (New England Biolabs) with the following thermal cycling conditions: 98°C for 2 minutes, 30 cycles of 20 seconds at 98°C, 20 seconds at 52°C, 15 seconds at 72°C, and a final extension at 72°C for 2 minutes. sgRNAs were *in vitro* transcribed using the HiScribe T7 High Yield RNA Synthesis Kit (New England Biolabs) according to manufacturer’s protocol. Transcription reactions were digested with 2 units RNase-free DNase I (New England Biolabs) at 37°C for 10 min; the reaction was stopped by adding EDTA to a final concentration of 35 mM and incubating at 75°C for 10 min. All guides were purified with RNAClean beads (Beckman Coulter) and quantified with the Quant-IT Ribogreen RNA Assay kit (ThermoFisher) according to the manufacturers’ protocols. Individual RNPs were prepared by incubating each sgRNA at 95°C for 2 minutes, then allowed to slowly come to room temperature over 5 minutes. Each sgRNA was then combined with its respective Cas9 in a 3:1 sgRNA:Cas9 molar ratio and incubated at 37°C for 10 minutes in cleavage reaction buffer (20 mM HEPES, pH 7.4, 150 mM KCl, 10 mM MgCl2, 5% glycerol). In 96-well format, 10 μg of gDNA was treated with 0.2 pmol, 0.8 pmol, 3.2 pmol, and 12.8 pmol of each RNP in 50 μL total volume in cleavage reaction buffer, in triplicate. Negative control reactions were assembled in parallel and did not include any RNP. gDNA was treated with RNPs for 4 hours at 37°C. Library preparation and sequencing were done according to protocols described previously [63] using the Illumina NextSeq platform, and ~3 million reads were obtained for each sample. Any SITE-Seq sites without off-target motifs located within 1 nt of the cut-site were considered false-positives and discarded.

### Targeted deep sequencing analysis

To measure indel frequencies, targeted deep sequencing analyses were done as previously described [65]. Briefly, we used two-step PCR amplification to produce DNA fragments for each on-target and off-target site. In the first step, we used locus-specific primers bearing universal overhangs with complementary ends to the TruSeq adaptor sequences (Supplemental Table 7). DNA was amplified with Phusion High Fidelity DNA Polymerase (New England Biolabs) using annealing temperatures of 60C, 64’C or 68C, depending on the primer pair. In the second step, the purified PCR products were amplified with a universal forward primer and an indexed reverse primer to reconstitute the TruSeq adaptors (Supplemental Table 7). Input DNA was PCR-amplified with Phusion High Fidelity DNA Polymerase (98°C, 15s; 61°C, 25s; 72°C, 18s; 9 cycles) and equal amounts of the products from each treatment group were mixed and run on a 2.5% agarose gel. Full-size products (~250bp in length) were gel-extracted. The purified library was deep sequenced using a paired-end 150bp MiSeq run.

MiSeq data analysis was performed using a suite of Unix-based software tools. First, the quality of paired-end sequencing reads (R1 and R2 fastq files) was assessed using FastQC (http://www.bioinformatics.babraham.ac.uk/projects/fastqc/). Raw paired-end reads were combined using paired end read merger (PEAR) [94] to generate single merged high-quality full-length reads. Reads were then filtered by quality [using Filter FASTQ [95]] to remove those with a mean PHRED quality score under 30 and a minimum per base score under 24. Each group of reads was then aligned to a corresponding reference sequence using BWA (version 0.7.5) and SAMtools (version 0.1.19). To determine indel frequency, size and distribution, all edited reads from each experimental replicate were combined and aligned, as described above. Indel types and frequencies were then cataloged in a text output format at each base using bam-readcount (https://github.com/genome/bam-readcount). For each treatment group, the average background indel frequencies (based on indel type, position and frequency) of the triplicate negative control group were subtracted to obtain the nuclease-dependent indel frequencies. Indels at each base were marked, summarized and plotted using GraphPad Prism. Deep sequencing data and the results of statistical tests are reported in Supplemental Table 3.

SITE-Seq cell-based validation was performed as previously described with minor modifications [63]. In brief, SITE-Seq sites were amplified from ~1,000-4,000 template copies per replicate and sequencing data from Cas9-treated samples were combined to minimize any variability due to uneven coverage across replicates. Cas9 cleavage sites were registered from the SITE-Seq data, and mutant reads were defined as any non-reference variant calls within 20 bp of the cut site. Sites with low sequencing coverage (< 1,000 reads in the combined, Cas9-treated samples or <200 reads in the reference samples) or >2% variant calls in the reference samples were discarded. Sites were tallied as cellular off-targets if they accumulated > 0.5% mutant reads in the combined, Cas9-treated samples. This threshold corresponded to sites that showed unambiguous editing when DNA repair patterns were visually inspected.

## List of Abbreviations

AAV: adeno-associated virus
BLESS: breaks labelling, enrichment on streptavidin and next-generation sequencing
BLISS: breaks labeling *in situ* and sequencing
bp: base pair
Cas: CRISPR-associated
Circle-seq: circularization for *in vitro* reporting of cleavage effects by sequencing
CjeCas9: *Campylobacter jejuni* Cas9
CMV: cytomegalovirus
CRISPR: clustered, regularly interspaced, short palindromic repeats
crRNAs: CRISPR RNAs
dCas9: “dead” Cas9
Digenome-seq: digested genome sequencing
DSB: double-strand breaks
dsODN: double-stranded oligodeoxynucleotide
DTS: dual target site
EF1α: elongation factor-1α
GeoCas9: *Geobacillus stearothemophilus*
GUIDE-seq: genome-wide unbiased identification of double strand breaks enabled by sequencing
HDR: homology-directed repair
HTGTS: high-throughput genome-wide translocation sequencing
IDLV: integrase-defective lentiviral vector
mESC: mouse embryonic stem cell
NHEJ: non-homologous end joining
NLS: nuclear localization signal
NmeCas9: *Neisseria meningitidis* (strain 8013) Cas9
NTS: NmeCas9 target site
PAM: protospacer adjacent motif
RNP: ribonucleoprotein
SauCas9: *Staphylococcus aureus* Cas9
sgRNA: single-guide RNA
SITE-Seq: selective enrichment and identification of tagged genomic DNA ends by sequencing
SpyCas9: *Streptococcus pyogenes* Cas9
T7E1: T7 Endonuclease 1
tracrRNA: *trans*-acting CRISPR RNA
tru-sgRNAs: truncated sgRNAs.

## Declarations

### Ethics approval and consent to participate

Not applicable.

### Consent for publication

Not applicable.

### Availability of data and material

The deep sequencing data from this study have been submitted to the NCBI Sequence Read Archive (SRA; http://www.ncbi.nlm.nih.gov/sra) under accession number XXXXXX. Plasmids will be made available via Addgene.

### Authors’ contributions

NA, XDG, PL, AE, AM, RI, AG, KES, and TW carried out genome editing experiments. TW and TGF carried out genotoxicity experiments and analyses. NA, XDG, PL, AM, AG, LJZ, and SAW carried out GUIDE-seq experiments and analysis. PDD, AHS, AML, KM, CKF, and PC carried out SITE-Seq experiments and analyses using NmeCas9 protein supplied by AM. XDG, PL, AM, LJZ, and SAW carried out additional bioinformatic and statistical analyses. All authors analyzed and interpreted data. NA, XDG, PC and EJS wrote the manuscript, and all authors edited the manuscript.

## Acknowledgments

We thank Yin Guo for technical assistance, Wen Xue and members of the Sontheimer and Wolfe laboratories for insightful comments and discussions, and Phil Zamore for the use of his flow cytometer. We are also grateful to the UMMS Deep Sequencing and Molecular Biology Core Laboratories for providing outstanding technical support services for this research project.

## References

1 Marraffini LA: CRISPR-Cas immunity in prokaryotes. Nature 2015, 526:55–61.

2 Mohanraju P, Makarova KS, Zetsche B, Zhang F, Koonin EV, van der Oost J: Diverse evolutionary roots and mechanistic variations of the CRISPR-Cas systems. Science 2016, 353:aad5147.

3 Sontheimer EJ, Barrangou R: The bacterial origins of the CRISPR genome-editing revolution. Hum Gene Ther 2015, 26:413–424.

4 Barrangou R, Fremaux C, Deveau H, Richards M, Boyaval P, Moineau S, Romero DA, Horvath P: CRISPR provides acquired resistance against viruses in prokaryotes. Science 2007, 315:1709–1712.

5 Brouns SJ, Jore MM, Lundgren M, Westra ER, Slijkhuis RJ, Snijders AP, Dickman MJ, Makarova KS, Koonin EV, van der Oost J: Small CRISPR RNAs guide antiviral defense in prokaryotes. Science 2008, 321:960–964.

6 Marraffini LA, Sontheimer EJ: CRISPR interference limits horizontal gene transfer in staphylococci by targeting DNA. Science 2008, 322:1843–1845.

7 Makarova KS, Wolf YI, Alkhnbashi OS, Costa F, Shah SA, Saunders SJ, Barrangou R, Brouns SJ, Charpentier E, Haft DH, et al: An updated evolutionary classification of CRISPR-Cas systems. Nat Rev Microbiol 2015, 13:722–736.

8 Deltcheva E, Chylinski K, Sharma CM, Gonzales K, Chao Y, Pirzada ZA, Eckert MR, Vogel J, Charpentier E: CRISPR RNA maturation by trans-encoded small RNA and host factor RNase III. Nature 2011, 471:602–607.

9 Garneau JE, Dupuis ME, Villion M, Romero DA, Barrangou R, Boyaval P, Fremaux C, Horvath P, Magadan AH, Moineau S: The CRISPR/Cas bacterial immune system cleaves bacteriophage and plasmid DNA. Nature 2010, 468:67–71.

10 Sapranauskas R, Gasiunas G, Fremaux C, Barrangou R, Horvath P, Siksnys V: The *Streptococcus thermophilus*CRISPR/Cas system provides immunity in *Escherichia coli*. Nucleic Acids Res 2011, 39:9275–9282.

11 Deveau H, Barrangou R, Garneau JE, Labonte J, Fremaux C, Boyaval P, Romero DA, Horvath P, Moineau S: Phage response to CRISPR-encoded resistance in Streptococcus thermophilus. J Bacteriol 2008, 190:1390–1400.

12 Mojica FJ, Diez-Villasenor C, Garcia-Martinez J, Almendros C: Short motif sequences determine the targets of the prokaryotic CRISPR defence system. Microbiol 2009, 155:733–740.

13 Gasiunas G, Barrangou R, Horvath P, Siksnys V: Cas9-crRNA ribonucleoprotein complex mediates specific DNA cleavage for adaptive immunity in bacteria. Proc Natl Acad Sci USA 2012, 109:E2579–2586.

14 Jinek M, Chylinski K, Fonfara I, Hauer M, Doudna JA, Charpentier E: A programmable dual-RNA-guided DNA endonuclease in adaptive bacterial immunity. Science 2012, 337:816–821.

15 Jiang W, Bikard D, Cox D, Zhang F, Marraffini LA: RNA-guided editing of bacterial genomes using CRISPR-Cas systems. Nat Biotechnol 2013, 31:233–239.

16 Cho SW, Kim S, Kim JM, Kim JS: Targeted genome engineering in human cells with the Cas9 RNA-guided endonuclease. Nat Biotechnol 2013, 31:230–232.

17 Cong L, Ran FA, Cox D, Lin S, Barretto R, Habib N, Hsu PD, Wu X, Jiang W, Marraffini LA, Zhang F: Multiplex genome engineering using CRISPR/Cas systems. Science 2013, 339:819–823.

18 Hwang WY, Fu Y, Reyon D, Maeder ML, Tsai SQ, Sander JD, Peterson RT, Yeh JR, Joung JK: Efficient genome editing in zebrafish using a CRISPR-Cas system. Nat Biotechnol 2013, 31:227–229.

19 Jinek M, East A, Cheng A, Lin S, Ma E, Doudna J: RNA-programmed genome editing in human cells. eLife 2013, 2:e00471.

20 Mali P, Yang L, Esvelt KM, Aach J, Guell M, DiCarlo JE, Norville JE, Church GM: RNA-guided human genome engineering via Cas9. Science 2013, 339:823–826.

21 Urnov FD: Genome Editing B.C. (Before CRISPR): Lasting Lessons from the “Old Testament”. CRISPR J 2018, 1:34–46.

22 Hsu PD, Lander ES, Zhang F: Development and applications of CRISPR-Cas9 for genome engineering. Cell 2014, 157:1262–1278.

23 Komor AC, Badran AH, Liu DR: CRISPR-Based Technologies for the Manipulation of Eukaryotic Genomes. Cell 2017, 168:20–36.

24 Sternberg SH, Doudna JA: Expanding the Biologist’s Toolkit with CRISPR-Cas9. Mol Cell 2015, 58:568–574.

25 Dominguez AA, Lim WA, Qi LS: Beyond editing: repurposing CRISPR-Cas9 for precision genome regulation and interrogation. Nat Rev Mol Cell Biol 2016, 17:5–15.

26 Wang H, La Russa M, Qi LS: CRISPR/Cas9 in Genome Editing and Beyond. Annu Rev Biochem 2016, 85:227–264.

27 Zetsche B, Gootenberg JS, Abudayyeh OO, Slaymaker IM, Makarova KS, Essletzbichler P, Volz SE, Joung J, van der Oost J, Regev A, et al: Cpf1 is a single RNA-guided endonuclease of a class 2 CRISPR-Cas system. Cell 2015, 163:759–771.

28 Shmakov S, Smargon A, Scott D, Cox D, Pyzocha N, Yan W, Abudayyeh OO, Gootenberg JS, Makarova KS, Wolf YI, et al: Diversity and evolution of class 2 CRISPR-Cas systems. Nat Rev Microbiol 2017, 15:169–182.

29 Fu Y, Foden JA, Khayter C, Maeder ML, Reyon D, Joung JK, Sander JD: High-frequency off-target mutagenesis induced by CRISPR-Cas nucleases in human cells. Nat Biotechnol 2013, 31:822–826.

30 Hsu PD, Scott DA, Weinstein JA, Ran FA, Konermann S, Agarwala V, Li Y, Fine EJ, Wu X, Shalem O, et al: DNA targeting specificity of RNA-guided Cas9 nucleases. Nat Biotechnol 2013, 31:827–832.

31 Bolukbasi MF, Gupta A, Wolfe SA: Creating and evaluating accurate CRISPR-Cas9 scalpels for genomic surgery. Nat Methods 2015, 13:41–50.

32 Casini A, Olivieri M, Petris G, Montagna C, Reginato G, Maule G, Lorenzin F, Prandi D, Romanel A, Demichelis F, et al: A highly specific SpCas9 variant is identified by *in vivo* screening in yeast. Nat Biotechnol 2018, 36:265–271.

33 Chen JS, Dagdas YS, Kleinstiver BP, Welch MM, Sousa AA, Harrington LB, Sternberg SH, Joung JK, Yildiz A, Doudna JA: Enhanced proofreading governs CRISPR-Cas9 targeting accuracy. Nature 2017.

34 Tsai SQ, Joung JK: Defining and improving the genome-wide specificities of CRISPRCas9 nucleases. Nat Rev Genet 2016, 17:300–312.

35 Tycko J, Myer VE, Hsu PD: Methods for optimizing CRISPR-Cas9 genome editing specificity. Mol Cell 2016, 63:355–370.

36 Yin H, Song CQ, Suresh S, Kwan SY, Wu Q, Walsh S, Ding J, Bogorad RL, Zhu LJ, Wolfe SA, et al: Partial DNA-guided Cas9 enables genome editing with reduced off-target activity. Nat Chem Biol 2018, 14:311–316.

37 Kleinstiver BP, Pattanayak V, Prew MS, Tsai SQ, Nguyen NT, Zheng Z, Joung JK: High-fidelity CRISPR-Cas9 nucleases with no detectable genome-wide off-target effects. Nature 2016, 529:490–495.

38 Slaymaker IM, Gao L, Zetsche B, Scott DA, Yan WX, Zhang F: Rationally engineered Cas9 nucleases with improved specificity. Science 2016, 351:84–88.

39 Jiang F, Taylor DW, Chen JS, Kornfeld JE, Zhou K, Thompson AJ, Nogales E, Doudna JA: Structures of a CRISPR-Cas9 R-loop complex primed for DNA cleavage. Science 2016, 351:867–871.

40 Jiang F, Zhou K, Ma L, Gressel S, Doudna JA: A Cas9-guide RNA complex preorganized for target DNA recognition. Science 2015, 348:1477–1481.

41 Jinek M, Jiang F, Taylor DW, Sternberg SH, Kaya E, Ma E, Anders C, Hauer M, Zhou K, Lin S, et al: Structures of Cas9 endonucleases reveal RNA-mediated conformational activation. Science 2014, 343:1247997.

42 Nishimasu H, Ran FA, Hsu PD, Konermann S, Shehata SI, Dohmae N, Ishitani R, Zhang F, Nureki O: Crystal structure of Cas9 in complex with guide RNA and target DNA. Cell 2014, 156:935–949.

43 Chylinski K, Makarova KS, Charpentier E, Koonin EV: Classification and evolution of type II CRISPR-Cas systems. Nucleic Acids Res 2014, 42:6091–6105.

44 Fonfara I, Le Rhun A, Chylinski K, Makarova KS, Lecrivain AL, Bzdrenga J, Koonin EV, Charpentier E: Phylogeny of Cas9 determines functional exchangeability of dual-RNA and Cas9 among orthologous type II CRISPR-Cas systems. Nucleic Acids Res 2014, 42:2577–2590.

45 Briner AE, Donohoue PD, Gomaa AA, Selle K, Slorach EM, Nye CH, Haurwitz RE, Beisel CL, May AP, Barrangou R: Guide RNA functional modules direct Cas9 activity and orthogonality. Mol Cell 2014, 56:333–339.

46 Esvelt KM, Mali P, Braff JL, Moosburner M, Yaung SJ, Church GM: Orthogonal Cas9 proteins for RNA-guided gene regulation and editing. Nat Methods 2013, 10:1116–1121.

47 Kim E, Koo T, Park SW, Kim D, Kim K, Cho HY, Song DW, Lee KJ, Jung MH, Kim S, et al: In vivo genome editing with a small Cas9 orthologue derived from Campylobacter jejuni. Nat Commun 2017, 8:14500.

48 Ran FA, Cong L, Yan WX, Scott DA, Gootenberg JS, Kriz AJ, Zetsche B, Shalem O, Wu X, Makarova KS, et al: In vivo genome editing using *Staphylococcus aureus* Cas9. Nature 2015, 520:186–191.

49 Mir A, Edraki A, Lee J, Sontheimer EJ: Type II-C CRISPR-Cas9 biology, mechanism and application. ACS Chemical Biology 2018:in press.

50 Zhang Y: The CRISPR-Cas9 system in Neisseria spp. Pathog Dis 2017, 75.

51 Zhang Y, Heidrich N, Ampattu BJ, Gunderson CW, Seifert HS, Schoen C, Vogel J, Sontheimer EJ: Processing-independent CRISPR RNAs limit natural transformation in Neisseria meningitidis. Mol Cell 2013, 50:488–503.

52 Zhang Y, Rajan R, Seifert HS, Mondragón A, Sontheimer EJ: DNase H activity of *Neisseria meningitidis* Cas9. Mol Cell 2015, 60:242–255.

53 Hou Z, Zhang Y, Propson NE, Howden SE, Chu LF, Sontheimer EJ, Thomson JA: Efficient genome engineering in human pluripotent stem cells using Cas9 from *Neisseria meningitidis*. Proc Natl Acad Sci USA 2013, 110:15644–15649.

54 Lee CM, Cradick TJ, Bao G: The *Neisseria meningitidis* CRISPR-Cas9 system enables specific genome editing in mammalian cells. Mol Ther 2016, 24:645–654.

55 Ma E, Harrington LB, O’Connell MR, Zhou K, Doudna JA: Single-Stranded DNA Cleavage by Divergent CRISPR-Cas9 Enzymes. Mol Cell 2015, 60:398–407.

56 Rousseau BA, Hou Z, Gramelspacher MJ, Zhang Y: Programmable RNA cleavage and recognition by a natural CRISPR-Cas9 system from *Neisseria meningitidis*. Molecular Cell 2018:in press.

57 Harrington LB, Doxzen KW, Ma E, Liu JJ, Knott GJ, Edraki A, Garcia B, Amrani N, Chen JS, Cofsky JC, et al: A Broad-Spectrum Inhibitor of CRISPR-Cas9. Cell 2017, 170:1224–1233 e1215.

58 Pawluk A, Amrani N, Zhang Y, Garcia B, Hidalgo-Reyes Y, Lee J, Edraki A, Shah M, Sontheimer EJ, Maxwell KL, Davidson AR: Naturally occurring off-switches for CRISPR-Cas9. Cell 2016.

59 Sontheimer EJ, Davidson AR: Inhibition of CRISPR-Cas systems by mobile genetic elements. Curr Opin Microbiol 2017, 37:120–127.

60 Rauch BJ, Silvis MR, Hultquist JF, Waters CS, McGregor MJ, Krogan NJ, Bondy-Denomy J: Inhibition of CRISPR-Cas9 with Bacteriophage Proteins. Cell 2017, 168:150–158 e110.

61 Hynes AP, Rousseau GM, Lemay M-L, Horvath P, Romero DA, Fremaux C, Moineau S: An anti-CRISPR from a virulent streptococcal phage inhibits *Streptococcus pyogenes* Cas9. Nature Microbiology 2017, 2:1374–1380.

62 Kearns NA, Pham H, Tabak B, Genga RM, Silverstein NJ, Garber M, Maehr R: Functional annotation of native enhancers with a Cas9-histone demethylase fusion. Nat Methods 2015, 12:401–403.

63 Tsai SQ, Zheng Z, Nguyen NT, Liebers M, Topkar VV, Thapar V, Wyvekens N, Khayter C, Iafrate AJ, Le LP, et al: GUIDE-seq enables genome-wide profiling of off-target cleavage by CRISPR-Cas nucleases. Nat Biotechnol 2014, 33:187–197.

64 Cameron P, Fuller CK, Donohoue PD, Jones BN, Thompson MS, Carter MM, Gradia S, Vidal B, Garner E, Slorach EM, et al: Mapping the genomic landscape of CRISPR-Cas9 cleavage. Nat Methods 2017, 14:600–606.

65 Bolukbasi MF, Gupta A, Oikemus S, Derr AG, Garber M, Brodsky MH, Zhu LJ, Wolfe SA: DNA-binding-domain fusions enhance the targeting range and precision of Cas9. Nat Methods 2015, 12:1150–1156.

66 Wilson KA, McEwen AE, Pruett-Miller SM, Zhang J, Kildebeck EJ, Porteus MH: Expanding the Repertoire of Target Sites for Zinc Finger Nuclease-mediated Genome Modification. Mol Ther Nucleic Acids 2013, 2:e88.

67 Guan C, Kumar S, Kucera R, Ewel A: Changing the enzymatic activity of T7 endonuclease by mutations at the beta-bridge site: alteration of substrate specificity profile and metal ion requirements by mutation distant from the catalytic domain. Biochemistry 2004, 43:4313–4322.

68 Cho SW, Kim S, Kim Y, Kweon J, Kim HS, Bae S, Kim JS: Analysis of off-target effects of CRISPR/Cas-derived RNA-guided endonucleases and nickases. Genome Res 2014, 24:132–141.

69 Fu Y, Sander JD, Reyon D, Cascio VM, Joung JK: Improving CRISPR-Cas nuclease specificity using truncated guide RNAs. Nat Biotechnol 2014, 32:279–284.

70 Hwang WY, Fu Y, Reyon D, Maeder ML, Kaini P, Sander JD, Joung JK, Peterson RT, Yeh JR: Heritable and precise zebrafish genome editing using a CRISPR-Cas system. PLoS One 2013, 8:e68708.

71 Ran FA, Hsu PD, Lin CY, Gootenberg JS, Konermann S, Trevino AE, Scott DA, Inoue A, Matoba S, Zhang Y, Zhang F: Double nicking by RNA-guided CRISPR Cas9 for enhanced genome editing specificity. Cell 2013, 154:1380–1389.

72 Santos-Pereira JM, Aguilera A: R loops: new modulators of genome dynamics and function. Nature Reviews Genetics 2015, 16:583–597.

73 Ciccia A, Elledge SJ: The DNA damage response: making it safe to play with knives. Molecular Cell 2010, 40:179–204.

74 Chen PB, Chen HV, Acharya D, Rando OJ, Fazzio TG: R loops regulate promoter-proximal chromatin architecture and cellular differentiation. Nature Structural & Molecular Biology 2015, 22:999–1007.

75 Aouida M, Eid A, Ali Z, Cradick T, Lee C, Deshmukh H, Atef A, AbuSamra D, Gadhoum SZ, Merzaban J, et al: Efficient fdCas9 Synthetic Endonuclease with Improved Specificity for Precise Genome Engineering. PLoS One 2015, 10:e0133373.

76 Zhu LJ, Holmes BR, Aronin N, Brodsky MH: CRISPRseek: a bioconductor package to identify target-specific guide RNAs for CRISPR-Cas9 genome-editing systems. PLoS One 2014, 9:e108424.

77 Tsai SQ, Nguyen NT, Malagon-Lopez J, Topkar VV, Aryee MJ, Joung JK: CIRCLE-seq: a highly sensitive in vitro screen for genome-wide CRISPR-Cas9 nuclease off-targets. Nat Methods 2017, 14:607–614.

78 Yan WX, Mirzazadeh R, Garnerone S, Scott D, Schneider MW, Kallas T, Custodio J, Wernersson E, Li Y, Gao L, et al: BLISS is a versatile and quantitative method for genome-wide profiling of DNA double-strand breaks. Nat Commun 2017, 8:15058.

79 Zhu LJ, Lawrence M, Gupta A, Pagés H, Kucukural A, Garber M, Wolfe SA: GUIDEseq: a bioconductor package to analyze GUIDE-Seq datasets for CRISPR-Cas nucleases. BMC Genomics 2017, 18:379.

80 Chen F, Ding X, Feng Y, Seebeck T, Jiang Y, Davis GD: Targeted activation of diverse CRISPR-Cas systems for mammalian genome editing via proximal CRISPR targeting. Nat Commun 2017, 8:14958.

81 Dong, Guo M, Wang S, Zhu Y, Wang S, Xiong Z, Yang J, Xu Z, Huang Z: Structural basis of CRISPR-SpyCas9 inhibition by an anti-CRISPR protein. Nature 2017, 546:436–439.

82 Shin CS, Jiang F, Liu J-J, Bray NL, Rauch BJ, Baik SH, Nogales E, Bondy-Denomy J, Corn JE, Doudna JA: Disabling Cas9 by an anti-CRISPR DNA mimic. Science Advances 2017:, in press.

83 Yang H, Patel DJ: Inhibition Mechanism of an Anti-CRISPR Suppressor AcrIIA4 Targeting SpyCas9. Mol Cell 2017.

84 Fu BX, Hansen LL, Artiles KL, Nonet ML, Fire AZ: Landscape of target:guide homology effects on Cas9-mediated cleavage. Nucleic Acids Res 2014, 42:13778–13787.

85 Pattanayak V, Lin S, Guilinger JP, Ma E, Doudna JA, Liu DR: High-throughput profiling of off-target DNA cleavage reveals RNA-programmed Cas9 nuclease specificity. Nat Biotechnol 2013, 31:839–843.

86 Kim S, Kim D, Cho SW, Kim J, Kim JS: Highly efficient RNA-guided genome editing in human cells via delivery of purified Cas9 ribonucleoproteins. Genome Res 2014, 24:1012–1019.

87 Zuris JA, Thompson DB, Shu Y, Guilinger JP, Bessen JL, Hu JH, Maeder ML, Joung JK, Chen Z-Y, Liu DR: Cationic lipid-mediated delivery of proteins enables efficient protein-based genome editing *in vitro* and *in vivo*. Nat Biotechnol 2015, 33:73–80.

88 Bisaria N, Jarmoskaite I, Herschlag D: Lessons from Enzyme Kinetics Reveal Specificity Principles for RNA-Guided Nucleases in RNA Interference and CRISPR-Based Genome Editing. Cell Syst 2017, 4:21–29.

89 Villefranc JA, Amigo J, Lawson ND: Gateway compatible vectors for analysis of gene function in the zebrafish. Dev Dyn 2007, 236:3077–3087.

90 Kearns NA, Pham H, Tabak B, Genga RM, Silverstein NJ, Garber M, Maehr R: Functional annotation of native enhancers with a Cas9-histone demethylase fusion. Nat Methods 2015, 12:401–403.

91 Mou H, Smith JL, Peng L, Yin H, Moore J, Zhang XO, Song CQ, Sheel A, Wu Q, Ozata DM, et al: CRISPR/Cas9-mediated genome editing induces exon skipping by alternative splicing or exon deletion. Genome Biol 2017, 18:108.

92 Gupta A, Hall VL, Kok FO, Shin M, McNulty JC, Lawson ND, Wolfe SA: Targeted chromosomal deletions and inversions in zebrafish. Genome Res 2013, 23:1008–1017.

93 Guschin DY, Waite AJ, Katibah GE, Miller JC, Holmes MC, Rebar EJ: A rapid and general assay for monitoring endogenous gene modification. Methods Mol Biol 2010, 649:247–256.

94 Zhang J, Kobert K, Flouri T, Stamatakis A: PEAR: a fast and accurate Illumina Paired-End reAd mergeR. Bioinformatics 2014, 30:614–620.

95 Blankenberg D, Gordon A, Von Kuster G, Coraor N, Taylor J, Nekrutenko A, Galaxy T: Manipulation of FASTQ data with Galaxy. Bioinformatics 2010, 26:1783–1785.

